# Systemic-to-mucosal trafficking of memory B cells contributes to humoral immunity in the upper respiratory tract

**DOI:** 10.64898/2026.04.17.719191

**Authors:** Eva Piano Mortari, Mattia Laffranchi, Bianca Laura Cinicola, Chiara Sugoni, Sabina Barresi, Valentina Marcellini, Emanuele Agolini, Christian Albano, Gabriele Volpe, Marco Scarsella, Ezio Giorda, Alessandro Sparaci, Reparata Rosa Di Prinzio, Salvatore Zaffina, Concetta Quintarelli, Cinzia Milito, Marco Anile, Isabella Quinti, Antonio Novelli, Ling Chen, Franco Locatelli, Silvano Sozzani, Rita Carsetti

## Abstract

Systemic vaccination induces serum antibodies and circulating memory B cells, protects from severe disease, but does not ensure sterilizing immunity in the upper respiratory tract, where many respiratory pathogens initiate infection. How systemic memory B cells contribute to mucosal immunity remains unclear. Using multiparametric flow cytometry, single-cell RNA and V(D)J sequencing, and functional analyses of paired blood and nasal/oropharyngeal samples, we characterized human B cells across systemic and mucosal compartments. Swab-derived B cells transcriptionally overlap with circulating activated memory B cells while exhibiting distinct features of activation, tissue retention, and spontaneous IgA/IgG secretion. Approximately 6% of mucosal B-cell clones were shared with blood, indicating systemic–mucosal connectivity. Both infection and vaccination expanded two circulating antigen-specific activated memory B cells subsets, whereas antigen-specific B cells accumulated in the upper respiratory tract only following local inflammation. The finding that B-cell recruitment is reactive rather than preemptive may explain the limited efficacy of parenteral vaccines and provides a rationale for developing integrated systemic–mucosal vaccination strategies.

## Introduction

The upper respiratory tract (URT) is the primary portal of entry for respiratory pathogens and a key battleground for controlling transmission^1–6^. Conventional vaccines induce the production of memory B cells (MBCs), plasmablasts, and plasma cells producing high-affinity antibodies. Despite their great effectiveness in reducing disease severity, hospitalisation rates, and mortality, available vaccines do not generate sterilising immunity^7,8^ and are unable to prevent infection and block viral transmission. In the case of SARS-CoV-2, the pandemic is no longer a global emergency ^9^, but the virus continues to evolve and generate highly infectious variants, responsible for the persistent circulation of the virus^4,7–9,11,12^.

Mucosal protection in the URT relies on a combination of locally produced secretory IgA (SIgA), serum-derived IgG, and tissue-resident memory B and T lymphocytes ^3,10–12^. SIgA is secreted by plasma cells that reside in the lamina propria and actively transported across epithelial barriers via the polymeric Ig receptor (pIgR)^5,13,14^. SIgA can be induced by intranasal vaccination, as demonstrated for SARS-CoV-2^15^ and SIgA generated against the Omicron variant show high neutralizing efficacy if compared to serum IgG^16,17^. Antibodies of IgG isotype derive from systemic immunity and reach mucosal sites by transudation from the serum, and a dedicated transporter, the neonatal Fc receptor, transfers IgG to the luminal surface of epithelial cells^18^. The local concentration of IgG is highest shortly after vaccination, coinciding with the peak level of IgG in the serum^7,8,19^. Nevertheless, IgG concentration in mucosal secretions is 100-fold lower than in serum^20^, reflecting a limited diffusion of IgG across mucosal surfaces of the URT. Therefore, locally produced IgA and serum IgG contribute to protecting mucosal surfaces.

Following parenteral vaccination, mucosal IgG levels transiently rise in parallel with serum titers^7,8,19,20^. Although systemic immunization induces robust serum antibody responses, it elicits minimal local immunity in the URT^4,21–23^. On the other hand, natural infection generates mucosal humoral responses^4^, including SIgA and tissue-resident memory B (BRM) and T (TRM)^14,24–28^. However, reinfections remain common, suggesting that also mucosal immunity in the URT cannot provide sterilizing immunity.

Although lung-resident BRM have been characterized in mice^4,14,26,27,29,30^, the presence, phenotype, and origin of their human counterparts in the nasal and oropharyngeal mucosa remain poorly understood. To prevent infection, MBCs must produce antibodies on the surface of the nasal and oropharyngeal mucosa, at the sites of viral entry.

Recent studies have identified various B cell types in swabs from the nasopharyngeal posterior wall and adenoids, and from the nasal middle turbinates^25^. CD69□ BRM-like cells were found in all swabs, but germinal center (GC) B cells were only detected in the posterior wall and adenoids swabs. After breakthrough infection, SARS-CoV-2-specific BRM were found in all posterior and adenoidal swabs but only in 31% of the medial turbinate samples^25^. These findings suggest that local antigen encounter promotes B-cell residency in the URT (in the tonsils and adenoids where GCs develop) but also underscore the immunological distinction between adjacent mucosal sites within the upper airways.

Here, we investigated whether systemic B cell immunity contributes to local protection in the URT.

By combining flow cytometry, single-cell RNA-seq, and clonal lineage analysis of B cells from nasal and oropharyngeal swabs and paired blood samples, we reveal a transcriptional and clonal continuum between circulating and swab-derived B cells. We show that swab-derived B cells exhibit phenotypic and transcriptional similarity to circulating activated MBCs (actMBCs).

Clonal and somatic hypermutation analyses reveal comparable mutation frequencies across circulating actMBCs and mucosal subsets and show that approximately 6% of swab-derived B cell clones are shared with blood, indicating exchange between compartments.

Together, these findings reveal a previously unrecognized pathway by which systemic immunity contributes to local protection of mucosal sites. Infection or vaccination drives the antigen-dependent expansion of actMBCs with mucosal-homing potential.

Recruitment to URT surfaces is, however, inflammation-dependent, suggesting that while conventional parenteral vaccination generates MBCs capable of mucosal migration, their recruitment is reactive rather than preemptive.

The low frequency of B cells in the swabs under steady-state conditions supports the notion that inflammatory signals may also regulate trafficking and retention of BRM cells, thus explaining why most respiratory viruses continue to circulate and cause disease.

## Results

### Phenotype of B cells in the upper respiratory tract

We analyzed B lymphocytes contained in the nasal and oropharyngeal swabs by flow-cytometry. Exfoliated epithelial cells and debris were excluded based on forward- and side-scatter properties (FSC-A vs SSC-A). CD45□SSC-A^+^ lymphocytes were gated, and CD19 expression was used to define the B-cell population (Fig. 1A). The frequency of lymphocytes was significantly lower in both nasal and oropharyngeal swabs compared to peripheral blood (p<0.0001) (Fig. 1B). B cells accounted for approximately 8-14% of lymphocytes in peripheral blood, but increased to ∼50% in the oropharyngeal swabs, while remaining low/rare in nasal swabs (Fig. 1C).

**Fig. 1:**
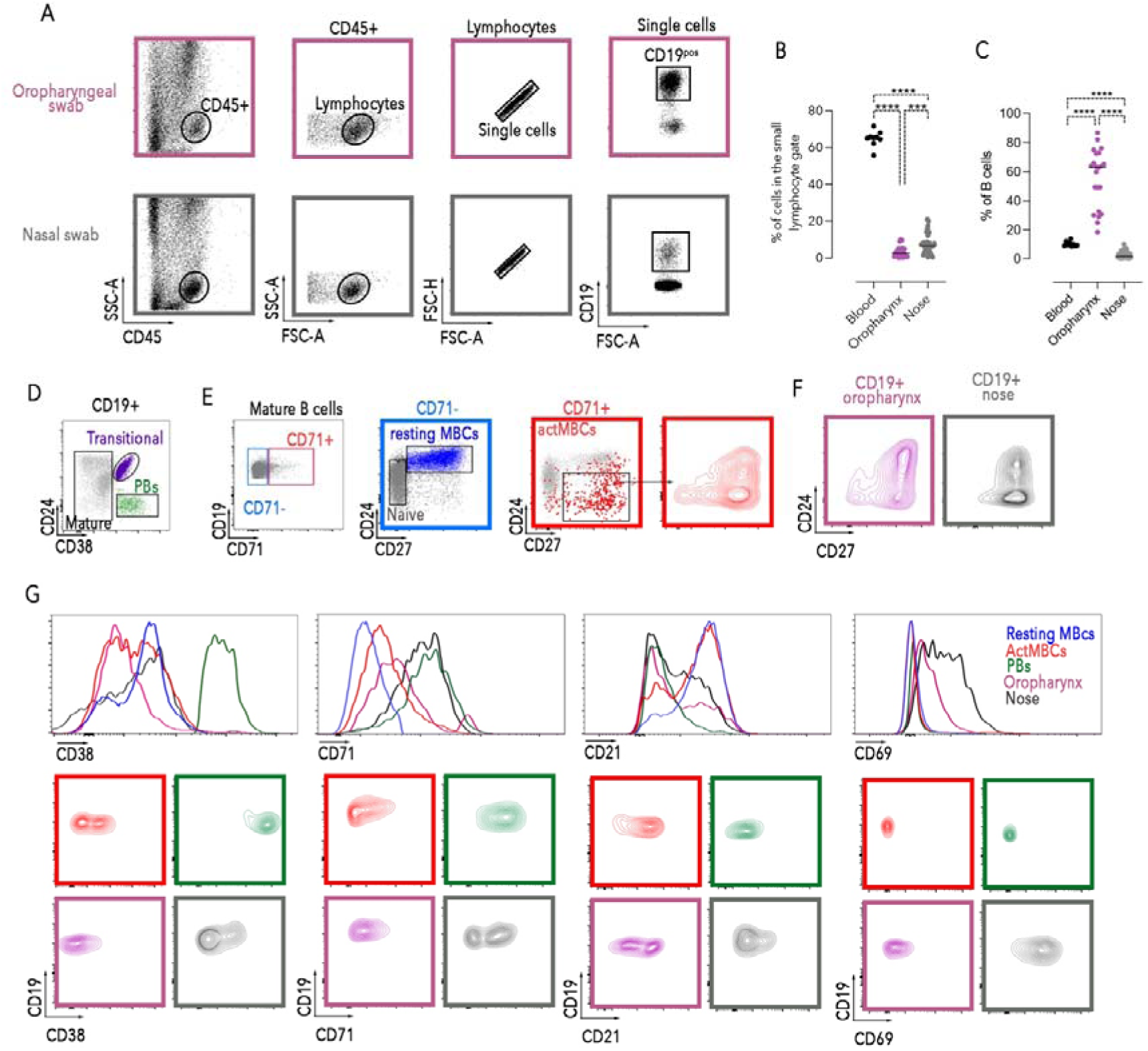
Phenotype of B cells isolated from nasal and oropharyngeal swabs. (A) FACS plots show the gating strategy to identify CD45^+^CD19^+^ cells in nasal and oropharyngeal swabs. (B-C) Graphs show the percentage of cells in the small lymphocyte gate and of CD19^+^ cells from peripheral blood, nasal (n=32) and oropharyngeal (n=24) swabs. (D) FACS plots show the gating strategies to identify B cell populations in adult peripheral blood with antibodies against CD19, CD24, CD27, CD38, CD21, and CD71. In the CD19+ B cell gate, transitional B cells (purple) are CD24^++^CD38^++^ and plasmablasts (PBs) (in green) are CD24^-^CD38^++^. (E) After the exclusion of plasmablasts (not-plasmablasts gate), we separate the B cells into CD71^-^ and CD71^+^. Among CD71^-^ we separate mature-naïve (CD24^+^CD27^-^, in grey) from the resting memory B cells (MBCs) population (in blue). CD71^+^ represent the activated MBCs (actMBCs in red, CD71^+^CD24^-/+^CD27^+^). (F) FACS plots of the expression of CD24 and CD27 in B cells isolated from oropharyngeal and nasal swabs. (G) Histogram and representative contour plots depict CD38, CD71, CD21 and CD69 expression, in resting MBCs, actMBCs and PBs isolated from blood, and B cells isolated from oropharyngeal and nasal swabs. Lines represent mean. The nonparametric Mann-Whitney t-test was used to evaluate statistical significance. The significance of the two-tailed P value is shown as ***p<0.001, ****p< 0.0001.

To further characterize the phenotype of B cells in the mucosal swabs, we performed detailed phenotypic profiling and compared swab-derived B cells with circulating B-cell subsets from the same individuals (Fig. 1D). In the peripheral blood, distinct B-cell populations were identified as previously described^31,32^. Transitional B cells were defined as CD24^++^CD38^++^, and plasmablasts (PBs) as CD24^-^CD38^++^ (Fig. 1D). Within mature B cells, CD71^+^ actMBCs^33,34^ were distinguished from CD71^-^ populations, which included CD24^+^CD27^-^naïve B cells and CD24^+^CD27^+^ resting MBCs (Fig. 1E).

B cells isolated from both nasal and oropharyngeal swabs displayed a phenotype closely resembling actMBCs, based on the expression of CD27, CD24 and CD38 (Fig. 1F and G). This interpretation was supported by their expression of CD71 and partial downregulation of CD21, consistent with an activated phenotype (Fig. 1G). CD24^-^CD38^++^ PBs were undetectable in the swabs. Furthermore, swab-derived B-cells exhibited higher levels of CD69, a marker associated with tissue residency and activation, distinguishing them from circulating peripheral B cells (Fig. 1G).

### scRNA-seq analysis defines two activated memory B-cell populations in the peripheral blood and a specific swab-derived B-cell subset

Given that our phenotypic analyses indicated that B cells recovered from nasal and oropharyngeal swabs were predominantly CD27^pos^, and to gain additional insight into the relationships between systemic and swab-derived B cells, we performed single-cell RNA sequencing (scRNA-seq) on resting MBCs, actMBCs, and PBs isolated from peripheral blood, alongside CD19^pos^ cells isolated from nasal and oropharyngeal swabs of four healthy donors.

We obtained 50.524 cells, which clustered into 12 transcriptionally distinct subsets organized into four major B-cell populations: resting MBCs (sorted as CD27^dull^ and CD27^bright^), actMBCs (CD27_Act), PBs, and swab-derived B cells (Fig. 2A). Clusters were annotated based on enrichment of the sorted populations and canonical B-cell marker expression (Fig. 2B and Suppl. Table 1).

**Fig. 2.**
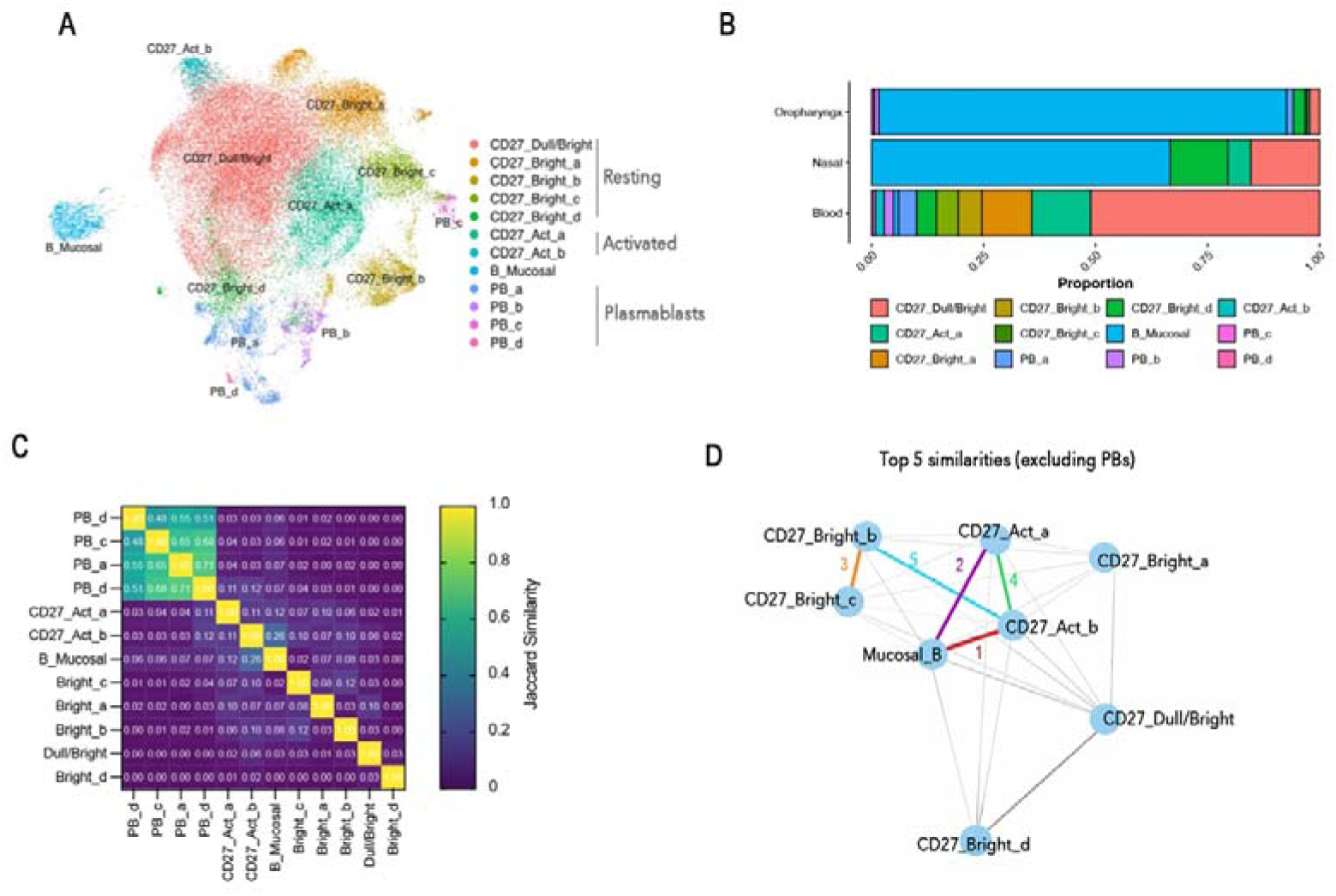
Mucosal B cells proximity with systemic activated memory B cells. (A) UMAP of 50,524 sorted B cells from blood, nasal, and oropharyngeal swabs from four healthy donors showing clustering of circulating B-cell subsets and swab-derived B cells (B_Mucosal). Each color represents a distinct cluster. (B) Stacked bar plot showing the proportional distribution of each cluster across the indicated tissue origins. (C) Jaccard similarity heatmap displaying overlap in marker gene expression among B-cell clusters. (D) Network plot showing the top five transcriptional similarities among B-cell clusters after exclusion of plasmablasts (PBs).

Among resting MBCs, we identified five clusters, including one shared CD27^dull/bright^ population and four clusters of CD27^bright^ MBCs (Fig. 2A and Suppl. Table 1).

The CD27^dull/bright^ cluster, containing most of the sorted CD27^dull^ cells, was enriched for the transcription factors *PAX5* and *BACH2* (Suppl. Table 1). *PAX5* maintains B-cell identity^35^, whereas *BACH2* promotes MBCs generation and survival and represses plasma cell differentiation by inhibiting *BLIMP*, either directly or in cooperation with *BCL6*^36–38^. In contrast, the CD27^bright^ MBC clusters displayed transcriptional profiles consistent with a more differentiated state^39^, characterized by genes linked to activation, protein synthesis, and energy metabolism (Suppl. Table 1).

PBs were identified by their transcriptomic signature, including *XBP1, IRF4*, *PRDM1*, *SDC1*, *CCR2*, *TRAM1*, *SEC61B,* and *KDEL*^33,40^, and showed diverse heavy- and light-chain expression profiles (Suppl. Table 1).

We quantified the relative abundance of each B-cluster in peripheral blood, nasal swab, and oropharyngeal swab samples (Fig. 2B). This analysis revealed clear tissue-specific enrichment patterns. In both nasal and oropharyngeal samples, the “B_Mucosal” cluster represented the predominant population (Fig. 2A and B), although smaller percentages of systemic B cells were also present (Fig. 2B). On the other hand, blood samples contained all identified clusters, reflecting the full spectrum of the CD27^+^ B-cell differentiation pathway. UMAP analysis highlighted two transcriptionally distinct clusters of actMBCs (“CD27_Act_a” and “CD27_Act_b”), both expressing the canonical actMBCs gene signature^33^ (Suppl. Table 1). To evaluate the transcriptional relationship across all clusters, we performed a Jaccard similarity analysis (Fig. 2C) followed by network visualization of the top inter-cluster similarities (excluding PBs - Fig. 2D). The Jaccard similarity heatmap showed that swab-derived B cells had a moderate overlap with systemic actMBCs subsets (Fig. 2C). Consistently, the network plot highlighted “CD27_Act_b” as the circulating subset with the strongest similarity to swab-derived B cells, followed by “CD27_Act_a” (Fig. 2 D) suggesting that swab-derived B cells share transcriptional programs with the two systemic actMBCs clusters while maintaining a distinct mucosal-specific identity.

To dissect the heterogeneity within actMBCs, we compared the transcriptomes of the two clusters identified by scRNA-seq. Differential gene expression analysis revealed marked transcriptional divergence between the two subsets (Fig. 3A). The “CD27_Act_a” cluster was enriched for *JCHAIN*, *CD38, PRKCB, ILR6,* and showed reduced expression of *PAX5,* suggesting a possible early commitment toward plasma cell differentiation^40–42^ (Fig. 3A and Suppl. Table 2). On the other hand, the “CD27_Act_b cells” expressed the surface markers *FCRL5, FCRL3, ITGAX* (CD11c), and *MS4A1* (CD20), while downregulating *CR2* (CD21) (Fig. 3A and Suppl. Table 2). The transcriptional profile of “CD27_Act_b” closely resembled that of effector MBCs (eMBC) FCRL5□ B cells induced by influenza infection^30,43^ or vaccination^44^, age-associated B cells (ABCs)^45,46^, and atypical CD21□ B cells^47^. Notably, the transcription factors T-bet and TCF-1 were also differently expressed by the two populations of actMBCs. The “CD27_Act_a” cluster expressed *TCF7* (TCF-1), and the “CD27_Act_b” expressed *TBX21* (T-bet).

**Fig. 3:**
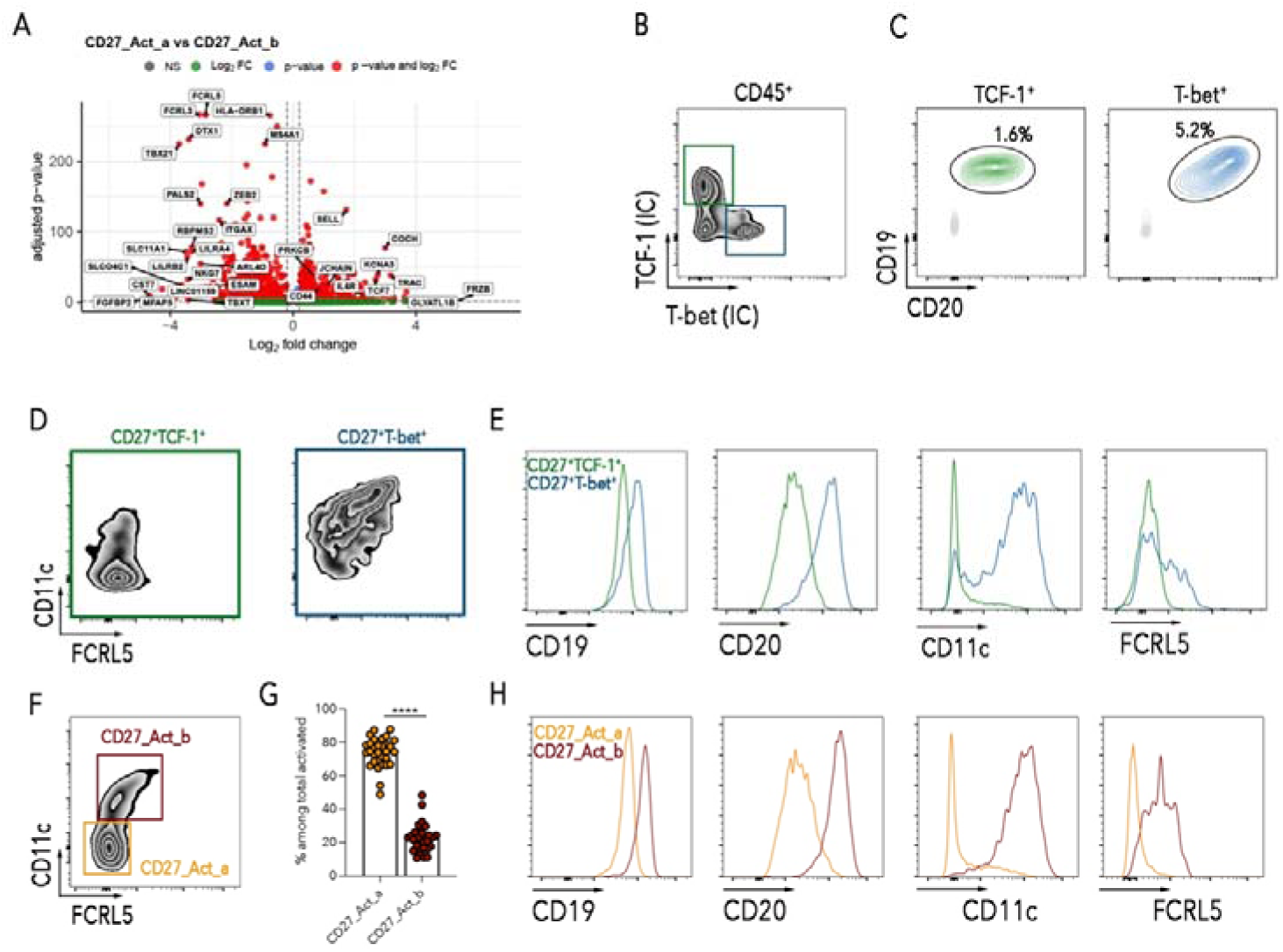
Characterization of activated MBCs in peripheral blood. (A) Volcano plot of differentially expressed genes between CD27_Act_a and CD27_Act_b clusters. (B) Zebra plots show the expression of T-bet and TCF-1 among CD45□ cells. (C) Contour plots identify CD19^+^CD20 B cells among T-bet□ and TCF-1□ populations. (D) Zebra plots of the expression of CD11c and FCRL5 among TCF-1□ and T-bet□ B cells. (E) Histogram overlay showing differential expression of CD19, CD20, CD11c, and FCRL5 between TCF-1□CD27^+^ and T-bet□CD27^+^ B cells subsets. (F) Representative zebra plots illustrating expression of FCRL5, CD11c, and T-bet in actMBCs to identify CD27_Act_a and CD27_Act_b subsets. (G) Column scatter graphs show the frequency of CD27_Act_a and CD27_Act_b among total actMBCs. (H) Expression of CD19, CD20, CD11c, and FCRL5 in CD27_Act_a and CD27_Act_b B cells. Columns depict mean + SEM. The nonparametric Wilcoxon matched pair t-test was used to evaluate statistical significance. The significance of the two-tailed P value is shown as ****p< 0.0001.

To validate these findings at the protein level, we analyzed TCF-1 and T-bet expression by flow cytometry. Within CD45□ cells, distinct TCF-1□ and T-bet□ populations were identified (Fig. 3B). We gated CD19□CD20□ B cells expressing each transcription factor to separate them from T and NK cells (Fig. 3C). The CD27□TCF-1□ subset was predominantly CD11c□FCRL5□, whereas the CD27□T-bet□ subset was CD11c□FCRL5□ (Fig. 3D). In agreement with the transcriptomic data, CD27□T-bet□ B cells displayed higher expression of CD19, CD20, CD11c, and FCRL5 compared to CD27□TCF-1□ B cells (Fig. 3E).

Based on these results, to simplify the staining procedure, we used the surface expression of FCRL5 and CD11c to identify CD27_Act_b and CD27_Act_a B cells (Fig. 3F), among CD71^+^ actMBCs. Both subsets were detectable in the peripheral blood of healthy adults, although CD27_Act_a cells were significantly more abundant (p<0.0001; Fig. 3G). Confirming our correct identification of the two populations, the mean fluorescence intensity of CD19, CD20, CD11c, and FCRL5 was significantly lower in CD27_Act_a than in CD27_Act_b cells, mirroring the protein expression pattern of CD27□TCF-1□ B cells (Fig. 3H).

Together, these results identify two distinct actMBC populations based on the expression of two transcription factors and two surface markers. The T-bet^+^ CD27_Act_b subset expresses FCRL5 and CD11c, whereas the TCF-1^+^ CD27_Act_a population is largely negative for CD11c and FCRL5. These two actMBCs populations differ in their transcriptional program and surface phenotype.

### Mucosal swab B cells comprise a mixture of CD27_Act_a and CD27_Act_b B cells

Considering the observation that swab-derived B cells share transcriptional programs with the two systemic clusters of actMBCs, we focused our subsequent analyses on the “B_Mucosal”, “CD27_Act_a”, and “CD27_Act_b” clusters (Fig. 4A).

**Fig. 4:**
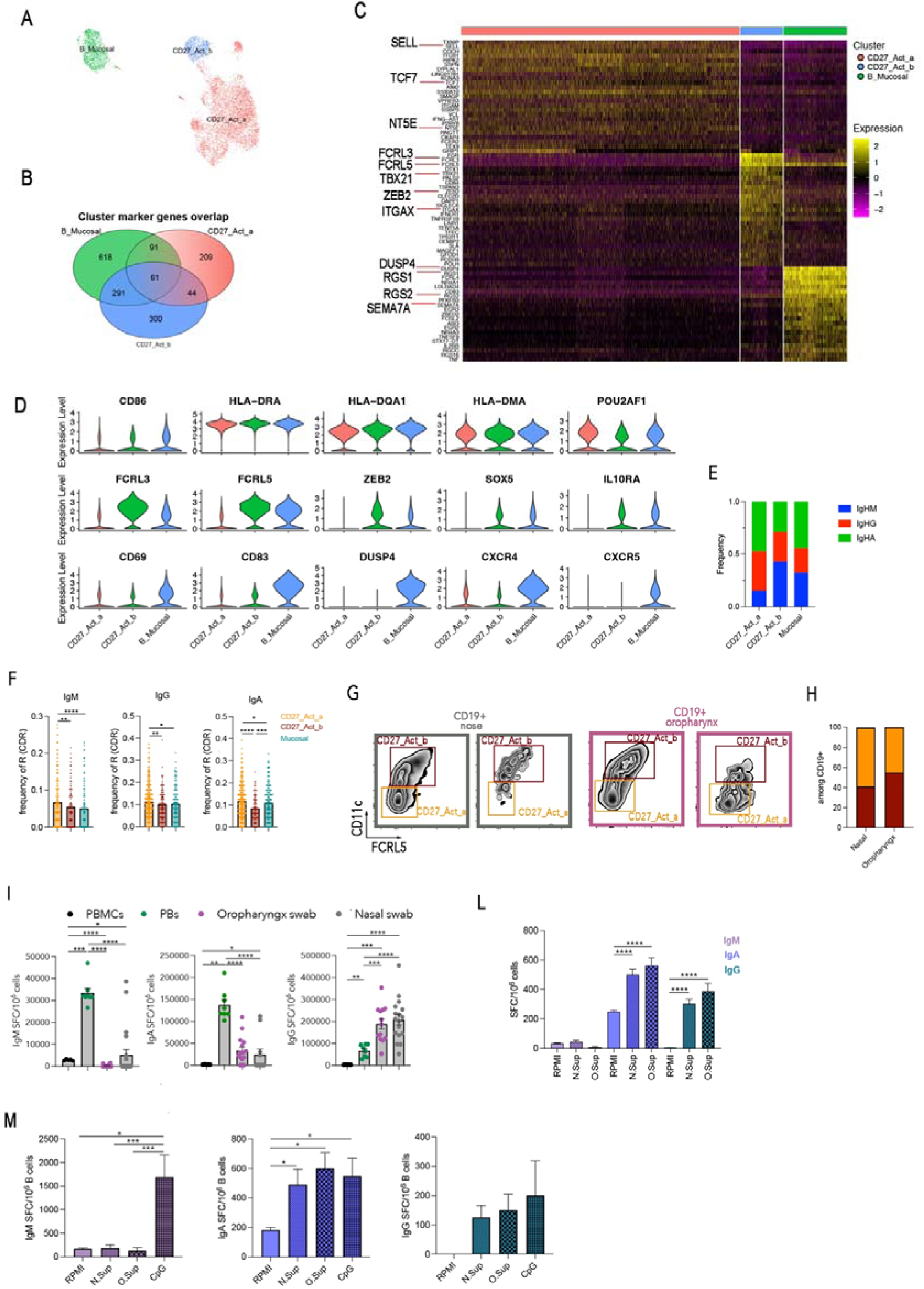
Characterization of swab-dervied B cells. (A) Reclustering of B_Mucosal, CD27_Act_a, and CD27_Act_b populations based on gene expression profiles. (B) Venn diagram showing overlap of marker genes between B_Mucosal, CD27_Act_a, and CD27_Act_b clusters. (C) Heat maps depicts the top 25 genes expressed by CD27_Act_a, CD27_Act_b and B_Mucosal clusters (D) Violin plots of selected genes highlighting shared and distinct transcriptional features among the three populations. (E) Stacked bar graph summarizing the frequency of each isotype within the indicated subsets. (F) Replacement mutation frequency in IgM□, IgA□, and IgG□ among CD27_Act_a, CD27_Act_b subsets, and swab-derived B cells. (G) Representative flow cytometry plots showing the distribution of CD27_Act_a and CD27_Act_b subsets in nasal and oropharyngeal compartments. (H) Frequency of CD27_Act_a and CD27_Act_b among total CD19□ B cells in nasal and oropharyngeal samples. (I) Graphs depict total IgA, IgG, and IgM spots per million cells among PBMCs, PBs, and swab-derived B cells. (L-M) Graphs depict the frequency of ASCs generated after the addition of nasal or oropharyngeal supernatants compared to RPMI alone in total PBMCs (L) or sorted activated MBCs (M). For sorted actMBCs stimulation with CpG was used as internal control. Columns depict mean + SEM. The nonparametric Wilcoxon pair signed rank test and Mann-Whitney t test were used to evaluate statistical significance. The significances of the two-tailed P value are shown as *p< 0.05, **p<0.01, ***p<0.001, ****p<0.0001.

Analysis of cluster marker overlap indicated that B cells derived from the swabs shared the greatest number of genes with “CD27_Act_b” (n=291) and a smaller set with “CD27_Act_a” (n=91) (Fig. 4B). The gene set enrichment analysis revealed that all three populations were enriched in pathways related to B-cell activation and migration, whereas the “B_Mucosal” and “CD27_Act_b” clusters also showed enrichment for pathways involved in B-cell differentiation, regulation of inflammatory and type II interferon responses (Suppl. Fig. 1). Comparative transcriptional profiling confirmed the shared activation programs among these subsets, with the strongest overlap in gene expression observed between “B_Mucosal” and “CD27_Act_b”. Heat maps of top 25 genes (Fig. 4C) and violin plots of representative genes (Fig. 4D) highlighted not only the proximity between these two clusters, but also partial similarity across all three clusters, indicating a continuum of activation states rather than fully segregated phenotypes. All three clusters expressed activation-associated transcripts such as *CD86*, *HLA-DRA*, *HLA-DQA1*, *HLA-DMA* and *POU2AF1*—the latter essential for MBC differentiation and GC function^46,48,49^ (Fig. 4D and Suppl. Table 1). *SELL* (CD62-L) which promotes B cell migration to the lymph nodes^50^ and *NT5E* (CD73), a marker often expressed by GC-derived MBCs^43^ were mostly expressed by “CD27_Act_a” (Fig. 4C). *FCRL3, FCRL5, ZEB2, SOX5, IL10RA* and *ITGAX,*were shared between “B_Mucosal” and “CD27_Act_b”, supporting their close transcriptional proximity (Fig. 4C and D). *ZEB2* has been identified as a central regulator of the T-bet^+^ B-cell program and sustains GC reactions during infection^51^. FCRL5 functions as an IgG receptor that enables immune complex sensing^52^, and can synergize with CD11c (*ITGAX*) by promoting phagocytosis and cell migration^53^, thereby supporting and enhancing early B-cell responses to infection^54^. *SOX5*, previously associated with tissue-resident B cells^25^, acts as a transcription factor that limits the proliferation of B cells during the GC reaction, ensuring the survival of cells expressing high-affinity antibodies and generation of PBs^55^. *FCRL3* plays a dual role: it enhances IL-10 production in B cells, thereby suppressing inflammatory cytokine secretion^56^, yet its ligation also amplifies TLR9-mediated B-cell proliferation, activation, and survival^57^. Interestingly, both “CD27_Act_b” and B cells derived from mucosal swabs expressed *IL10RA* suggesting a potential role in immune regulation and resolution of inflammation^58^ which is particularly relevant in the upper airway where excessive or prolonged inflammation can compromise airways integrity and breathing.

At the same time, swab-derived B cells expressed unique markers, including *CD69* (early activation marker) and *CD83* (late activation marker) which are involved in tissue residency^25,59^, as well as *DUSP4*, a regulator of activation^60^. Swab B cells also expressed chemokine receptors such as *CCR1*, *CXCR4* and *CXCR5* all involved in mucosal immunity and acting as tissue retention molecules^61–63^ (Fig. 4D and Suppl. Table 1).

*RGS1* and *RGS2*, members of the R4 regulator of G protein signaling (RGS) family, known to modulate the activation of B cells, attenuate their response to chemokines^64,65^, and limit their movement out of tissues^66,67^, were also enriched in the “B_Mucosal” cluster (Fig. 4C). V(D)J sequencing (scVDJ-seq) analysis confirmed that actMBCs and swab B cells are antigen-experienced populations. The “CD27_Act_a” cluster predominantly expressed class-switched isotypes, whereas the “CD27_Act_b” cluster retained substantial IgM expression. The “B_Mucosal” cluster exhibited an intermediate composition, reflecting the relative contributions of both actMBC subsets (Fig. 4E). Mutation frequency of swab-derived B cells mirrored that of “CD27_Act_b” for IgM and IgG sequences but resembled “CD27_Act_a” for IgA (Fig. 4F). Together, these transcriptional features indicate that swab-derived B cells are activated, tissue-retained, and with immunoregulatory functions, thus occupying a niche that is related but also molecularly distinguishable from systemic actMBCs.

To evaluate the similarity between swab-derived B cells and actMBCs at the protein level, we analyzed the expression of CD11c and FCRL5 (Fig. 4G). Both actMBCs populations were detected at mucosal sites, confirming that swab-derived B cells encompass a mixture of CD27_Act_a and CD27_Act_b subsets (Fig. 4H).

As antibodies represent the B cell effector molecules mediating protection against infection, we evaluated the immunoglobulin-secreting capacity of B cells isolated from the nasal and oropharyngeal swabs. Spontaneous antibody production was assessed because B cells recovered from swabs derived from a non-sterile environment and cannot be cultured under sterile conditions. ELISpot assays were performed on B cells isolated from the swabs and compared to sorted peripheral blood PBs, CD71^+^ actMBCs and CD71^-^ resting MBCs. Total PBMCs stimulated *in vitro* for 5 days were used as positive control. Swab-derived B cells secreted substantial amount of IgG, and at lower extent IgA (Fig. 4I and Suppl. Fig. 2A). IgM secretion was generally low across all mucosal samples. As expected, PBs secreted high levels of antibodies of all isotypes, whereas sorted actMBCs and resting MBCs did not secrete immunoglobulin in the absence of stimulation (Suppl. Fig. 2A). These data indicate that B cells recovered from the nasal and oropharyngeal mucosa are functionally active and poised to secrete antibodies, indicating that they contribute to local IgA and IgG production, although they do not have either the transcriptome or the phenotype of PBs.

ActMBCs have been described as immediate precursors of antibody-secreting cells (ASCs)^33^. We compared the response of actMBCs and resting MBCs to either a T-independent (TI) or T-dependent (TD) stimulus. CpG ODN-2006, a synthetic oligonucleotide that binds to TLR9, was chosen as a representative TI stimulus; whereas the TD signal was mimicked by engaging CD40 and Ig on B cells. Both stimulations were performed in the presence of IL-4 and IL-2^68^.

We sorted resting MBCs and actMBCs and stimulated them for 3 days. Non-B cells (sorted as CD19^-^ cells) were used as feeder layer (Suppl. Fig. 2B). We found that, after 3 days, in response to a TD or TI stimulus, actMBCs produced significantly more switched ASCs than resting MBCs (Suppl. Fig. 2B), thus confirming that actMBCs are more prone to differentiate into ASCs than resting MBCs.

We reasoned that the ability of swab-derived B cells to secrete antibodies without stimulation may be explained by their previous exposure to activating signals within the URT microenvironment, probably including microbiota-derived factors. To directly address whether soluble factors present in the URT are sufficient to promote B-cell activation and differentiation, we cultured PBMCs for 24 hours in RPMI alone or supplemented with cell-free supernatants from nasal or oropharyngeal swabs. All cellular components were removed from swab samples before culture. Exposure to nasal- or oropharyngeal-derived supernatants resulted in the increased frequency of ASCs, mostly of switched isotype, compared to RPMI alone (Fig. 4L). To investigate whether actMBCs have the intrinsic ability to generate such a rapid response, we show that sorted actMBCs, cultured alone, respond to nasal- or oropharyngeal-derived supernatants by increasing the frequency of switched ASCs within 24 hours (Fig. 4M). These results demonstrate that soluble mediators present in the mucosal environment are sufficient to drive B-cell functional activation.

Collectively, these data demonstrate that swab-derived B cells represent a specialized population that is transcriptionally related to systemic actMBCs but adapted for local tissue residency. It remains to be determined whether swab-derived B cells only reflect resident B cell-immunity or include a systemic MBCs component migrated to the mucosa.

### Kinetics of CD27_Act_a and CD27_Act_b in response to infection or vaccination

To investigate the role of the two populations of actMBCs *in vivo*, we used the response to SARS-CoV-2 vaccination or infection as a working model. In a cohort of health care workers (HCWs), we measured the frequency of Spike-specific B cells 9 months after the 2^nd^ dose of the BNT162b2 mRNA vaccine (T270 n=11), and 10 days after the 3^rd^ dose (n=11). We also analyzed the peripheral blood of HCWs who experienced a breakthrough infection (INF; n=15, 10 days after infection). After vaccination, B cells that recognize the Spike protein can be detected in all CD27^+^ populations (Suppl. Fig. 3A) including CD27^dull^ and CD27^bright^ resting MBCs. In response to recent stimulation (vaccination or infection), antigen-specific B cells increase significantly among total actMBCs (p=0.05 after 3^rd^ dose) and PBs (p=0.015; Suppl. Fig. 3A). Infection generated more Spike-specific actMBCs than vaccination (p=0.013; Suppl. Fig. 3A).

In another cohort of HCWs analyzed before and 10 days after vaccination or after infection (T270 and after 3^rd^ dose n=12; INF=7), we dissected the actMBCs into CD27_Act_a and CD27_Act_b and evaluate the frequency of antigen-specific cells (Fig. 5A). Nine months after the 2^nd^ dose, Spike-specific CD27_Act_a were significantly higher than Spike-specific CD27_Act_b (p=0.034; Fig. 5A). Administration of the 3^rd^ dose or infection significantly increased the frequency of Spike-specific CD27_Act_b (3^rd^ p=0.001 and INF p=0.0005) and PBs (p=0.002 and p=0.0005) compared to the T270 (Fig. 5A).

**Fig. 5:**
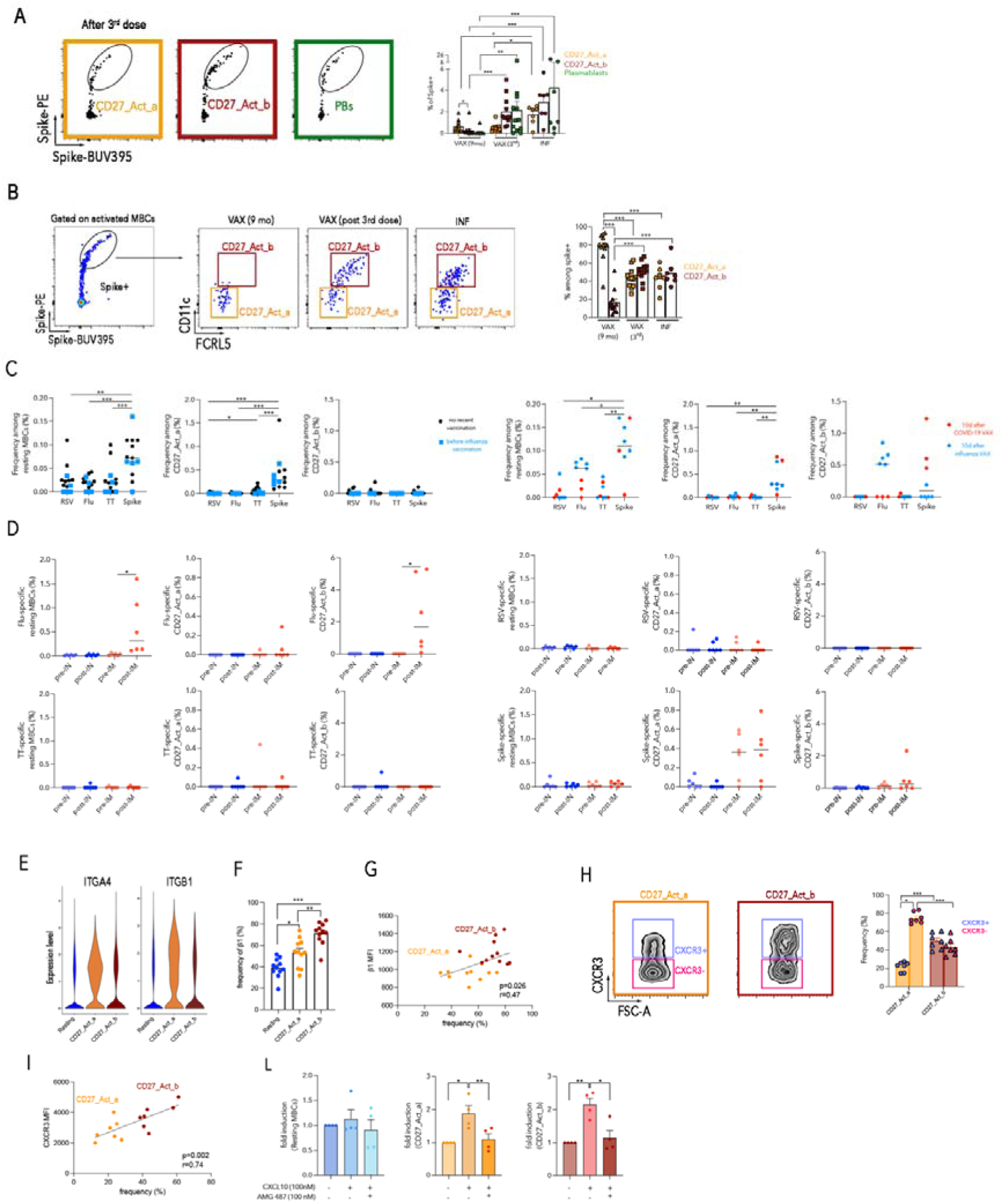
Antigen-specific MBCs: (A) Gating strategy to identify Spike-specific CD27_act_a, CD27_Act_b and PBs. Frequency of Spike-specific B cells CD27_Act_a, CD27_Act_b and PBs among the same subjects analyzed 9 months after the 2^nd^ dose (n=12), 10 days after the 3^rd^ dose (n=7) after a breakthrough infection. (B) Gating of Spike-specific B cells among actMBCs. Representative FACS plots of the expression of CD11c and FCRL5 among Spike-specific actMBCs 9 months after the 2^nd^ dose, 10 days after the 3^rd^ dose or infection. Column scatter graph depicts the percentage of CD27_Act_a and CD27_Act_b among Spike-specific actMBCs. (C) Graphs depict the frequencies of RSV-, Flu-, TT- and SARS-CoV-2-specific B cells among resting, CD27_Act_a and CD27_Act_b MBCs. In black samples that have been not recently vaccinated (n=8). In blue (square) samples before the influenza vaccination (n=5; not recently vaccinated for other antigen). In red samples that have been vaccinated with COVID-19 mRNA vaccine 10 days before (n=3), and in blue (circle) samples that received influenza vaccination 10 days before (n=5, same as square samples). (E) Graphs show the frequencies in children of RSV-, Flu-, TT- and SARS-CoV-2-specific B cells among resting, CD27_Act_a and CD27_Act_b MBCs. In blue samples that received intranasal vaccination (n=6) and in red samples that have been vaccinated with intramuscular vaccine (n=6) (F) Violin plots of the expression level of ITGA4 and ITGB1. (G) Frequency of β1□ resting MBCs, CD27_Act_a and CD27_Act_b. (H) Bivariate analysis of β1 expression, plotting β1 mean fluorescence intensity (MFI) against the frequency of β1□ cells for CD27_Act_a and CD27_Act_b subsets. Each dot represents a donor. (I) Representative zebra plots illustrating CXCR3^-^ and CXCR3^+^ among CD27_Act_a and CD27_Act_b subsets. Column scatter graph illustrates the frequency of CXCR3^+^ and CXCR3^-^ among CD27_Act_a and CD27_Act_b subsets. (L) Bivariate analysis of CXCR3 expression, plotting CXCR3 mean fluorescence intensity (MFI) against the frequency of CXCR3□ cells for CD27_Act_a and CD27_Act_b subsets. Each dot represents a donor. (M) Bar graphs display the fold induction of migration relative to spontaneous migration (set to 1) for resting, CD27_Act_a and CD27_Act_b MBCs (n=4). Columns depict mean + SEM (individual data points shown). The nonparametric Mann-Whitney t-test, the nonparametric Wilcoxon matched pair signed-rank test and the parametric paired t-test were used to evaluate statistical significance. The significance of the two-tailed P value is shown as *p< 0.05, **p<0.01, ***p< 0.001, ****p< 0.0001. Linear regression analysis was used to assess the relationship between (MFI) and the frequency (%). The dashed line indicates the linear regression fit. Pearson correlation analysis was performed to quantify the strength of the association. Statistical significance was defined as p < 0.05.

We investigated the relative contribution of CD27_Act_a and CD27_Act_b to the Spike-specific actMBC pool (Fig. 5B). We found that 9 months after the 2^nd^ dose, 80% of the Spike-specific actMBCs belonged to the CD27_Act_a population (Fig. 5B). Antigen-specific CD27_Act_b increased significantly after vaccination (p=0.0005) or infection (p=0.0007) when they represented 50% of the activated pool (Fig. 5B).

These data indicate that the CD27_Act_b subset strongly expands in an antigen-specific manner early after a recent stimulus (vaccination or infection). This population is transiently present in the peripheral blood, whereas the CD27_Act_a B cells appear to be continuously generated^69,70^.

To confirm the dependence of actMBCs on a recent antigen exposure, we analyzed the frequency of B cells with specificities other than Spike in the resting and actMBC populations. We quantified MBCs specific for the respiratory syncytial virus (RSV) fusion glycoprotein, the influenza hemagglutinin from H1N1/Victoria/2022 (Flu), and tetanus toxoid (TT), as representative antigens from pathogens or vaccines encountered in the past (Suppl. Fig 3B). Samples were collected during a period when RSV and influenza virus did not circulate. In addition, none of the subjects had recently received the diphtheria-pertussis-tetanus vaccine, and the last SARS-CoV-2 mRNA vaccine dose had been administered four years earlier. Five samples were analyzed before the influenza vaccination (blue squares; Fig. 5C).

We found that resting MBCs specific for RSV, influenza (Flu), and TT were detectable in all samples (Fig. 5C). The frequency of resting MBCs specific for Spike was significantly higher, likely reflecting the cumulative effect of repeated mRNA vaccine doses (Fig. 5C). Among CD27_Act_a MBCs, only those specific for Spike were detectable.

We also analyzed samples from individuals recently vaccinated against influenza (n=5) or SARS-CoV-2 (n=3). Flu-specific resting MBCs and CD27_Act_b B cells were higher in those samples that received influenza vaccination (Fig. 5C, in blue) and Spike-specific CD27_Act_b cell increased after the COVID-19 booster dose (Fig. 5C, in red). For samples analyzed before (blue squares) and after influenza vaccination (blue circles) the increased of flu-specific B cells was significant in the resting MBCs (p=0.005) and in the CD27_Act_b (p=0.004).

Together, these findings confirm the interpretation that CD27_Act_b B cells represent a short-lived population induced by a recent stimulation.

We extended these findings to a pediatric cohort that received either the intranasal (IN) or intramuscular (IM) influenza vaccine. IM had a significant impact specifically on flu-specific B cells: after IM vaccination there was a significant increase in both resting MBCs (p=0.031) and CD27_Act_b B cells (p=0.031) specific for influenza (Fig. 5E), while after IN vaccination we did not observe an increase in flu-specific B cells in peripheral blood. The frequencies of MBCs specific for the other antigens tested (RSV, TT, and Spike) did not change in the pediatric cohort (Fig. 5D), consistent with the fact that these subjects were specifically stimulated with the influenza antigen through IM vaccination. These results are consistent with the early generation of CD27_Act_b B cells after antigen stimulation. Moreover, they suggest that stimulation of the URT alone is not sufficient to generate systemic immunological memory.

We investigated whether CD27_Act_a and CD27_Act_b B cells are equipped for migration to the URT. Migration to the URT is guided by VLA-4 integrin^17^, composed of integrin α4 (ITGA4) and integrin β1 (ITGB1), which binds to vascular cell adhesion molecule-1 (VCAM-1)^71^. *ITGA4* and *ITGB1* expression was higher in actMBCs, especially in the “CD27_Act_a” cluster, compared to resting MBCs (Fig. 5E). At the protein level, significantly more CD27_Act_b cells were β1^+^ cells compared to CD27_Act_a (p=0.0029, Fig. 5F) B cells and β1 mean fluorescence intensity (MFI) positively correlated with β1^+^ cell frequency (p=0.026, Fig. 5G).

Recruitment of B cells to the airways requires CXCR3^29,59,72^, a chemokine receptor also necessary for protection against infections^63,73^. We directly evaluated the expression of CXCR3 by flow cytometry. We found that 20% of CD27_Act_a were CXCR3□, whereas 50% of CD27_Act_b expressed CXCR3 (Fig. 5H), demonstrating a significantly higher frequency of CXCR3 expression among CD27_Act_b compared with CD27_Act_a (p=0.0006). Among CXCR3□ cells, CXCR3 MFI was also higher in CD27_Act_b than in CD27_Act_a (Fig. 5I). Notably, CXCR3 frequency and CXCR3 MFI showed a strong and significant positive correlation (p=0.002; r = 0.74, Fig. 5I), indicating that samples with higher proportions of CXCR3□ cells also displayed increased receptor expression levels.

To validate the function of CXCR3-expression, we assessed the migratory response of B cell subsets to CXCL10, a canonical CXCR3 ligand. CXCL10 selectively induced the migration of total actMBCs (p=0.029), including both CD27_Act_a (p=0.033) to CD27_Act_b cells (p=0.009) (Fig. 5L). In contrast, total B cells, naïve B cells and resting MBCs did not exhibit significant migration in response to CXCL10, (Fig. 5L and Suppl. Fig. 3C). Pre-treatment with the selective CXCR3 antagonist AMG 487 abolished CXCL10-driven migration, confirming the specificity of the response (Fig. 5L). Notably, CD27_Act_b cells, exhibited a slightly higher response to CXCL10 than CD27_Act_a, consistent with their higher degree of membrane receptor expression. Collectively, these findings demonstrate that CXCR3 is functionally active in actMBCs, likely underpinning their preferential recruitment to inflamed airway tissues.

These data suggest that actMBCs express the correct equipment for trafficking to URT tissues and respond to the inflammatory chemokine CXCL10.

### Antigen-specific B cells in the upper respiratory tract

To define the conditions that promote B cell homing to the surface of the URT, we analyzed nasal and oropharyngeal swabs from a few individuals belonging to the vaccinated HCWs cohort described above. Swabs were collected in three different states: (1) after recovery from SARS-CoV-2 infection, (2) during acute inflammation of the URT (cold-like symptoms and SARS-CoV-2 PCR-negative), and (3) in the absence of inflammation (healthy, asymptomatic individuals). Spike-specific B cells were detected in all nasal swabs and in one oropharyngeal swab in individuals recovering from COVID-19 (Fig. 6A). In contrast, Spike-specific B cells were absent from both nasal and oropharyngeal swabs in healthy, non-inflamed individuals, indicating that homing of antigen-specific B cells to the URT does not occur constitutively in the absence of inflammation. Interestingly, during symptomatic SARS-CoV-2-negative URT inflammation, a low but detectable frequency (∼0.8%) of Spike-specific B cells was observed in nasal and oropharyngeal swabs, suggesting that inflammation alone is sufficient to promote B-cell migration to the mucosa (Fig. 6A). Recovery of viable CD45^+^ lymphocytes was less efficient in oropharyngeal swabs than nasal samples (Fig. 6B), explaining our fewer data on antigen-specific B cells in that compartment.

**Fig. 6:**
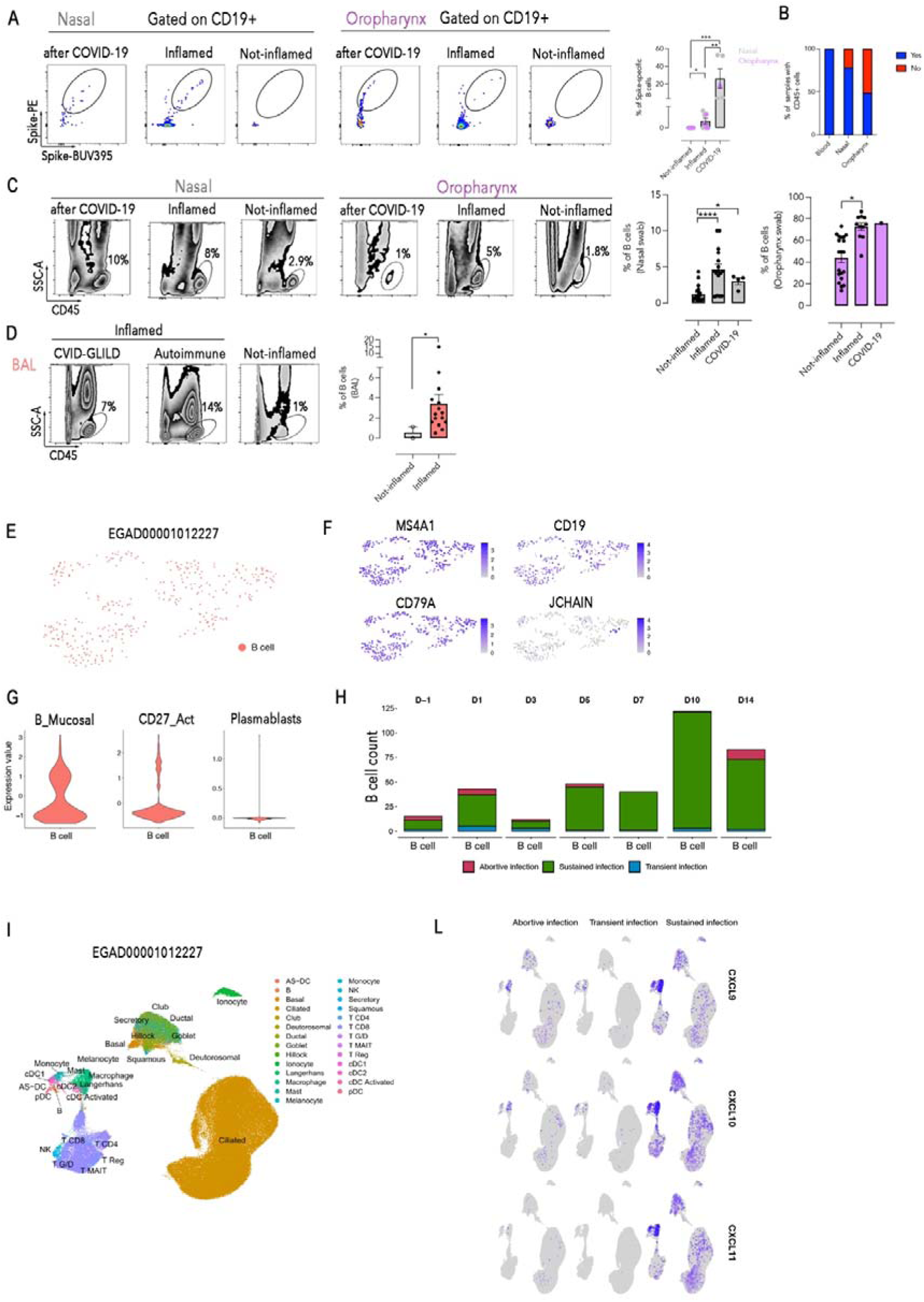
B cells in URT and in BAL. (A) Representative FACS plots and frequency of Spike-specific B cells in nasal and oropharyngeal swabs from healthcare workers (HCWs) collected in three conditions: following recovery from COVID-19 (n=4), during symptomatic but SARS-CoV-2–negative upper respiratory inflammation (n=7), and during asymptomatic, non-inflamed states (n=9). Spike-specific B cells were abundant post-infection, detected at low levels during inflammation, and absent in the non-inflamed state. (B) Bars represent the percentage of recovery success of viable lymphocytes from peripheral blood, and nasal and oropharyngeal swabs. (C) Representative zebra FACS plot of CD45^+^ cells and frequency of total CD19□ B cells in nasal (left panel) and oropharyngeal (right panel) swabs across the three conditions (Nasal swabs: COVID-19 n=4, Inflammed n=16, Not-Inflammed n=22; Oropharyngeal swabs: COVID-19 n=1, Inflammed n=11, Not-Inflammed n=20). Inflammation was associated with a significant increase in total B cells compared to the non-inflamed state. (D) Representative zebra FACS plots of CD45^+^ and frequency of CD19^+^ B cells in BAL samples from patients with GLILD, autoimmune disease and non-inflamed disease (tumor suspicion). (E) UMAP of 363 B cells nasopharyngeal swabs of healthy volunteers infected with SARS-CoV-2 from Lindeboom et al., 2024. (F) Feature plot showing the expression of indicated genes in the previous UMAP space. (G) Violin plots of selected gene signatures inferred from our dataset (Mucosal_B, CD27 actMBCs, and plasmablasts). (H) Stacked bar plot showing the absolute counts of B cells distribution stratified by the volunteer’s response to infection across the follow up. (I) UMAP of the entire dataset of nasopharyngeal swabs from healthy volunteers infected with SARS-CoV-2 from Lindeboom et al., 2024. Each color represents a distinct cluster. (L) Feature plot showing the expression of indicated genes in the previous UMAP space stratified by response to infection. Columns depict mean + SEM. The nonparametric Mann-Whitney t-test and the nonparametric Wilcoxon matched pair signed-rank test were used to evaluate statistical significance. The significance of the two-tailed P value is shown as *p< 0.05, **p<0.01, ***p< 0.001, ****p< 0.0001.

Quantification of CD45^+^ and CD19□ B cells revealed a significantly increased frequency in both nasal and oropharyngeal swabs during inflammation compared to the non-inflamed state, supporting a generalized B cell influx in response to mucosal inflammation (Fig. 6C). To determine whether inflammation-driven lymphocyte recruitment extends to the lower respiratory tract (LTR), we analyzed bronchoalveolar lavage (BAL) samples from patients with inflammatory lung conditions, including Common Variable Immunodeficiency (CVID)-associated granulomatous-lymphocytic interstitial lung disease (GLILD), suspected autoimmune disease and viral lung infection, as well from patients with suspected lung tumor (non-inflamed controls). Recovery of CD45^+^ lymphocytes and of CD19^+^ B cells was significantly higher in BAL samples from patients with inflammatory disease (Fig. 6D), indicating that mucosal inflammation promotes lymphocytes accumulation across anatomically distinct compartments.

Collectively, these data suggest that inflammation is a prerequisite for B cell recruitment to the URT and LRT, and that the presence of cognate antigen further enhances the local accumulation of antigen-specific B cells.

We show that the immune response to vaccination or infection generates circulating actMBCs with the potential ability to migrate to the respiratory tract. Local inflammatory cues, however, are indispensable to enable the recruitment, and antigen retention at the mucosa further promotes their persistence or expansion.

In order to further support our hypothesis, we re-analyzed a publicly available scRNA-seq dataset (Lindeboom et al., 2024, EGAD00001012227)^74^ from single-cell multi-omics profiling of nasopharyngeal swabs of participants in a controlled human infection study in which seronegative volunteers were intranasally infected with SARS-CoV-2. The infection resulted in three outcomes: abortive, transient, and sustained infection. After extracting B cells and applying quality filtering, due to the limited cell number (363 cells) we decided to analyze the B population as a whole (Fig. 6E). Still, we confirmed their purity by expression of canonical B-cell lineage genes (MS4A1, CD19, CD79A) (Fig. 6F). The plasmablast marker JCHAIN was absent (Fig. 6F). To further support this, we inferred our B cell clusters gene signatures, and we obtained an overlap with Mucosal_B cells, an enrichment of actMBCs but not with plasmablast (Fig. 6G). The co-existence of these two signatures, mucosal residency combined with an activated MBCs profile, suggests that these B cells have been actively recruited during the course of the infection, in line with our proposed model of inflammation-driven URT B-cell homing.

When stratifying total B-cell counts by day of sampling and infection outcome, we observed that at day -1 B cells were almost absent from the nasopharynx swabs in all conditions. A progressive increase in B-cell numbers was observed only in the sustained infection group, becoming most prominent from day 5 onward and peaking around day 10 (Fig. 6H). In contrast, B cells were almost undetectable in individuals with transient or abortive infection. In addition, when probing the expression of the chemokines recognized by CXCR3 (CXCL9, CXCL10 and CXCL11), we could observe their strong upregulation in the myeloid compartment from the sustained infection group, both in terms of average expression and percentages of cells positive (Fig. 6L). A lower expression was also observed in the epithelial compartment. The transient and abortive infection groups showed a restricted expression of CXCL9, CXCL10 and CXCL11 in dendritic cells. To summarize, in transiently infected individuals, the innate immune response was sufficient to control and clear the virus rapidly^74^, before the recruitment of adaptive B-cell populations becomes necessary. On the other hand, in sustained infection individuals, in which the SARS-CoV-2 virus was not promptly cleared by the innate immune system, there is a substantial recruitment of B cells and an upregulation of chemokines that recruits CXCR3^+^ cells.

Together, these findings converge on a model in which local inflammation, rather than antigen alone, allows B cell entry into the respiratory mucosa, with cognate antigen subsequently consent the accumulation of the recruited antigen-specific pool.

### Clonal structure, isotype distribution, and shared clone in peripheral blood and swab-derived B cells

To demonstrate whether systemic B cells contribute to B cell immunity in the URT, we performed scVDJ-seq to characterize the clonal architecture of B cells across the blood, nasal and oropharyngeal compartments. Clonal families were defined based on paired CDR3 sequences, and lineage trees were reconstructed using the Immcantation package^75,76^ (Suppl. Data).

A total of 36.692 unique clones corresponding to 40.969 single B cells were identified (Fig. 7A). Most of the clones were singlets, although expanded clones (≥6 cells) were evident, particularly in blood-derived samples. Isotype analysis revealed that IgM dominated both small and large clones, while class-switched isotypes such as IgG1, IgG2 and IgA were more frequent in singlets or small clones, and IgE^+^ B cells were absent (Fig. 7B-left panel). Isotype usage varied across tissues, with IgM enriched in the blood, IgA1 preferentially expressed in swab-derived B cells, and IgA2 and IgG2 significantly more represented in oropharyngeal B cells. Nasal-derived B cells exhibited increased IgD usage.

**Fig. 7:**
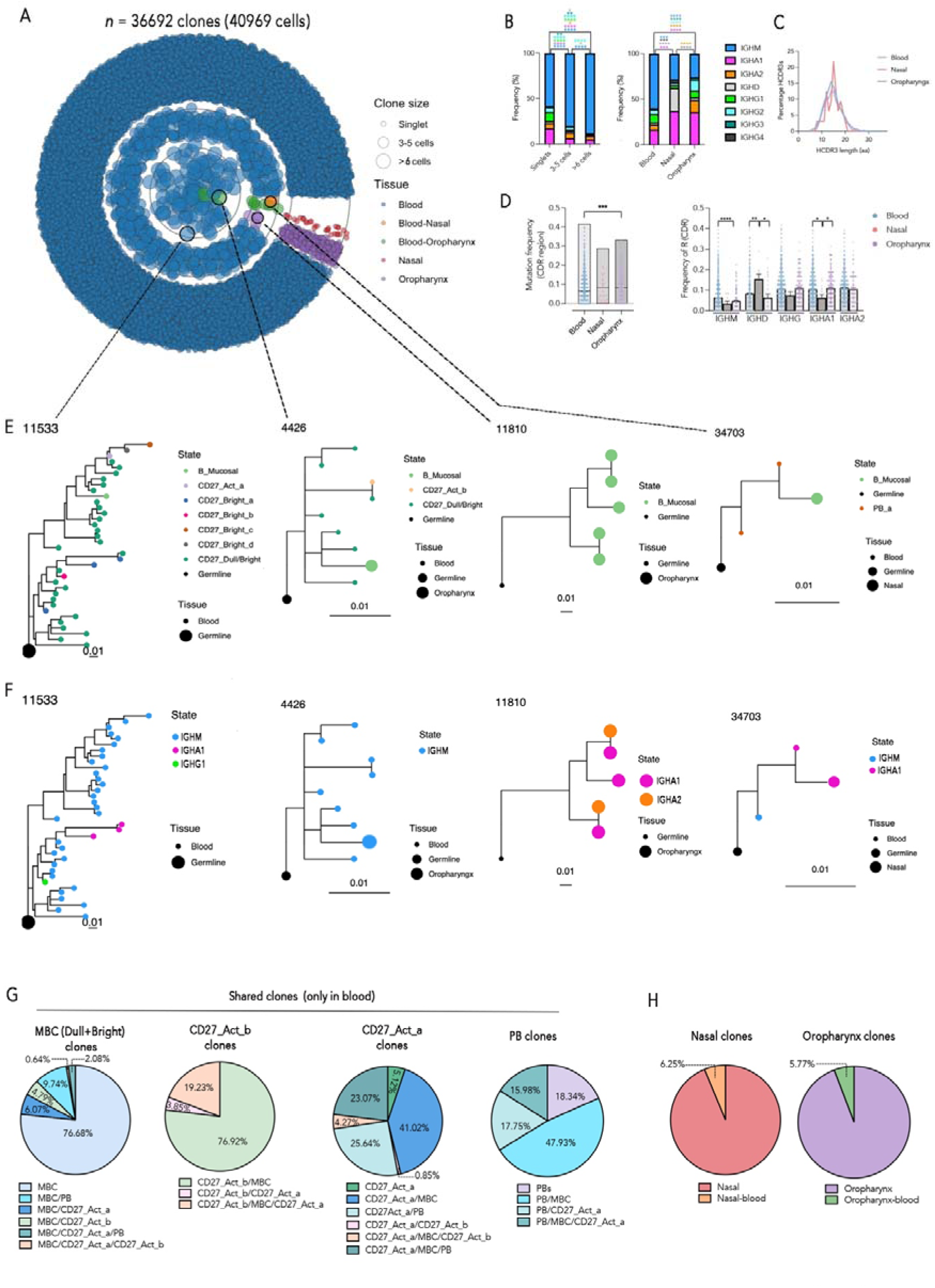
Clonal structure, isotype distribution, and shared clones in peripheral blood and swab-derived B cells. (A) Circular plot of clonal architecture across blood, nasal, and oropharyngeal samples (n = 36,692 clones, 40,969 single B cells). Each dot represents a single B cell; dot size corresponds to clone size; color indicates tissue of origin. Representative expanded clones with inter-compartment relationships are highlighted. (B) Isotype distribution across clones of increasing size (left panel) and by tissue origin (right panel). (C) HCDR3 length distribution (amino acids) across blood, nasal, and oropharyngeal B cells. (D) Mean of replacement somatic mutation frequency in VH CDR regions by tissue of origin (left panel) and stratified by isotype (right panel). (E) Representative clonal tree of expanded clones showing the distribution of B cell states (CD27-defined subsets and swab B cells) across blood and mucosal compartments. (F) Corresponding clonal phylogenies annotated by isotype. (G) Composition of blood-derived expanded clones. Left: clones containing MBCs (CD27^dull^ and CD27^bright^). Middle: clones containing CD27_Act_b and CD27_Act_a cells. Right: clones containing plasmablasts. Pie charts indicate relative contributions of B cell subsets within each clonal type. (H) Composition of swab-derived expanded clones from nasal and oropharyngeal samples. Pie charts show the proportion of tissue-restricted clones versus clones shared with blood. Pairwise statistical comparisons of immunoglobulin isotype distributions between cell groups were performed using a chi-square test of independence. Significance levels were indicated as follows: *p < 0.05, **p < 0.01, ***p < 0.001, ****p < 0.0001.

Analysis of HCDR3 length distributions revealed no substantial differences across compartments (Fig. 7C). Mutation frequency of non-ambiguous amino acids mismatches in the VH CDRs was slightly higher in oropharyngeal B cells (Fig. 7D-left panel) although this trend was not clearly apparent when stratifying by isotype (Fig. 7D-right panel) and might be due to the higher frequency of IgM MBCs in the blood. The mutation frequency was lowest in IgM MBCs as reported before^39,77^. Nasal IgD B cells were strongly mutated confirming previous results^78^ (Fig. 7D-right panel).

Expanded clones were identified in both blood and mucosal compartments (Fig. 7A). Lineage trees of representative expanded clones revealed both blood-restricted and mixed blood-mucosa clones (Fig. 7E), suggesting trafficking between compartments or a local diversification from common progenitors. Large blood-restricted clones remained primarily IgM+ (e.g., clone 11533) (Fig. 7F).

To further resolve the composition of expanded blood-derived clones, we analyzed the composition of the clonal trees (Fig. 7G). Clones containing resting MBCs were largely restricted to MBCs (76.68%), with minor contributions from “CD27_Act_a” (6.07%), “CD27_Act_b” (4.79%), or PBs (9.74%). “CD27_Act_b” clones showed strong clonal ties to MBCs (76.92%) and in part with “CD27_Act_a” (19.23%). PB-containing clones often included MBCs (47.93%) and/or “CD27_Act_a”, while a smaller fraction (18.34%) consisted exclusively of PBs.

Finally, clones from swab-derived samples were largely tissue-restricted, although approximately 6% were shared with circulating blood B cells (Fig. 7H). These findings highlight a complex B-cell landscape in the upper respiratory mucosa, composed of both compartmentalized and systemically connected clonal populations and indicate that the contribution of the blood to mucosal sites is at least 6%.

## Discussion

The SARS-CoV-2 pandemic has led to the extraordinary effort to vaccinate most of the world population. Vaccination protects from severe disease but does not prevent infection. The virus continues to circulate and generate variants. MBCs induced by mRNA vaccines are detected in the peripheral blood of all vaccinated individuals and effectively respond to breakthrough infections by proliferating, differentiating into PBs and producing antibodies in the serum and at mucosal sites^4,7,8,22,79–81^.

Spike-specific MBCs have been identified in the spleen^82^ and lymphoid tissues of vaccinated organ donors^83^, independently of prior SARS-CoV-2 infection^28^. This observation supports the notion that MBCs disseminate broadly after their initial induction, establishing systemic immune surveillance. However, Spike-specific MBCs were only found in the lungs of 31% of vaccinated-uninfected donors, and in 62.5% of vaccinated-infected donors^28^. Similarly, Spike-specific MBCs were in all adenoidal and tonsillar swabs, but in only 31% of the nasal swabs following breakthrough infection^25^. These findings highlight a discrepancy between the continuous presence of antigen-specific MBCs in lymphoid tissues and blood and their transient representation at mucosal surfaces—precisely where immediate protection is most critical.

Our study identifies the mechanism regulating the contribution of systemic B cells to mucosal immunity in the URT (Fig. 8).

**Fig. 8:**
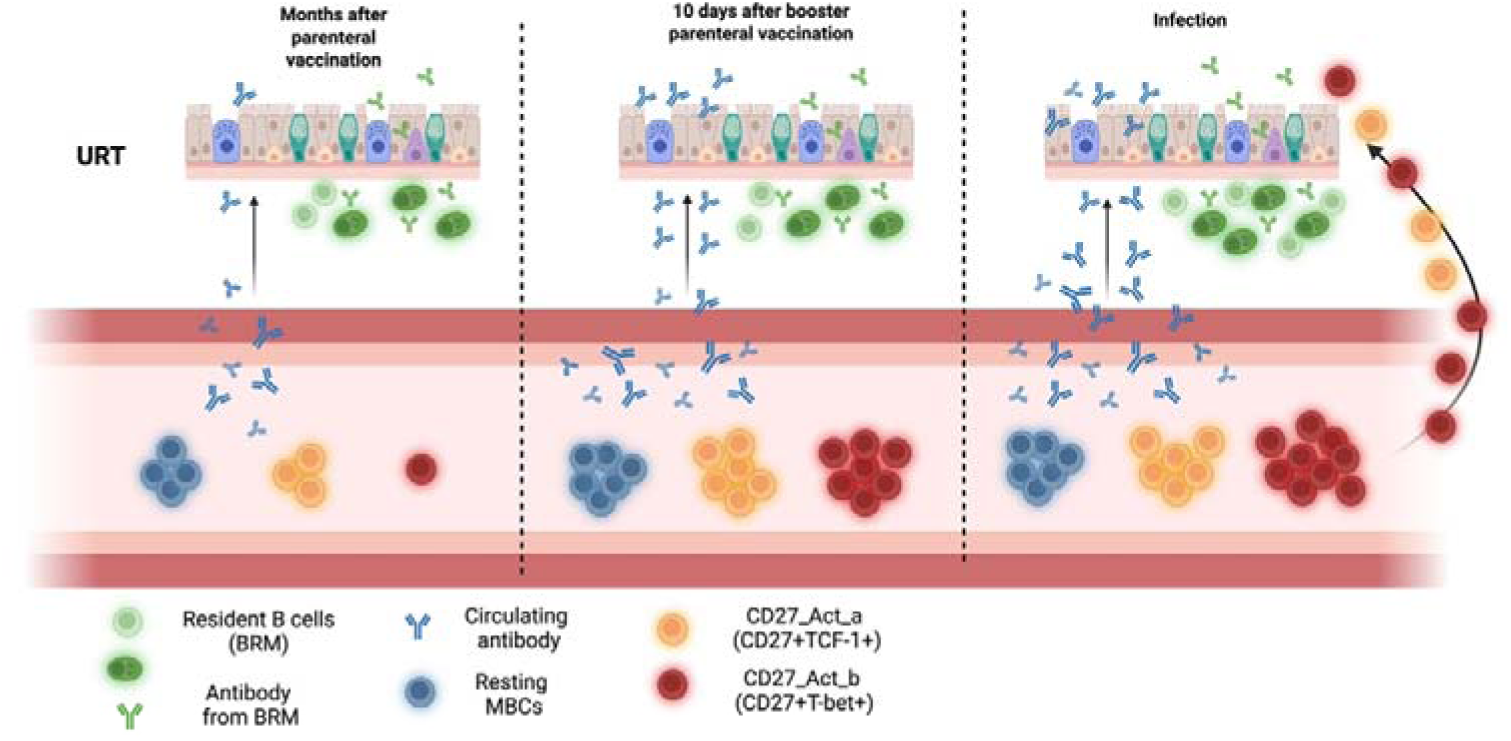
Schematic representation of main findings.

Using flow-cytometry, single-cell RNA and V(D)J sequencing of paired blood and nasal/oropharyngeal samples, we found that B cells isolated from nasal and oropharyngeal swabs share marker expression and transcriptional features with two populations of circulating actMBCs. Taking SARS-CoV-2 as a model antigen, we studied the in vivo kinetics of actMBCs. We show that CD27_Act_a B cells remain detectable months after immunization, whereas the CD27_Act_b population transiently^84^ expands after vaccination or infection.

The transcriptomic signature, surface marker phenotype, expression of T-bet, and strong dependence on recent stimulation all suggest that CD27_Act_b B cells may correspond to MBCs of extrafollicular origin^85^. The presence within this population of MBCs expressing highly mutated IgG and IgA, as well as less mutated IgM, suggests that both pre-existing class-switched and IgM MBCs may participate in the extrafollicular response and subsequently acquire additional functional competence. CD27_Act_b B cells may be related to the CD27^-^ populations that have been described as atypical and double negative MBC^45,46,86^. In contrast, the “CD27_Act_a” subset, based on its transcriptomic signature, surface phenotype, and kinetic profile, may correspond to MBCs generated in the GC. In the context of SARS-CoV-2 infection, GCs have been shown to persist for several months^69^, which may explain the sustained detection of antigen-specific, highly mutated, class-switched CD27_Act_a MBCs in peripheral blood. The expression of TCF-1 is consistent with a GC origin^87^. Further studies are warranted to determine whether TCF-1 also contributes to the biology of human B-1–like cells^88^. Notably, CD5 expression was not detected in either CD27_Act_a or CD27_Act_b cells (data not shown).

The transcriptome of swab B cells strongly overlaps with “CD27_Act_b”, although by flow cytometry we show that also CD27_Act_a are found in the swabs. At the same time, swab-derived B cells express unique markers involved in tissue residency^25,59^ and retention^61–63^.

### Our question was whether systemic memory B cells migrate to the nasal and oropharyngeal surfaces and contribute to the local defense

Based on the limited number of cells recovered from the swabs, we found that 6% of these B cells belonged to clones also present in blood, providing evidence for systemic-to-mucosal connectivity. Given the sparse cellularity of swab samples and the dominance of distinct clones within individual tissues reported in a repertoire study^83^, this value should be interpreted as a conservative lower bound rather than an absolute estimate of the extent of this phenomenon. We do not know whether the frequency of shared clones increases during infection or inflammation. In the samples analyzed here, the majority of swab-derived B cells belonged to clones not detected in the circulation, consistent with the presence of locally not circulating tissue-resident memory B cell populations that may acquire an activated phenotype in situ.

The nasal mucosa is uniquely enriched in IgD-producing B cells, a feature also observed in the oropharynx and tonsils^78,89,90^. In fact, a substantial fraction of plasma cells in nasal-associated lymphoid tissue express IgD^91^, which increases to 60% in individuals with selective IgA deficiency^92^, suggesting a specialized role for IgD in nasal immune defense.

Interestingly, both nasal and oropharyngeal mucosal tissues produce more IgA1 than peripheral blood reflecting a shared reliance on IgA-mediated protection at the mucosal surfaces of the URT. IgA2 was rarely expressed by swab B cells, but it is the predominant antibody produced by mucosal resident plasma cells. In the mucosa of the gut, MBCs are exposed to the combined effects of bacterial products and the cytokines APRIL and TGF-β, promoting switching to IgA2^93,94^. Similarly, in the tonsils, the expression of APRIL is polarized to the subepithelial space where it may induce class switching and promote plasma cells survival^95^. The different microenvironment of the swabs may explain why B cells continue to express IgG or IgA1 and do not switch to IgA2.

Total B cells and SARS-CoV-2 specific B cells were found at high frequencies in the swabs after SARS-CoV-2 infection. During non-COVID-19-related inflammation, the frequency of total B cells was comparable to COVID-19 infection, while the Spike-specific B cells were present at significantly lower frequency. This observation suggests a dual requirement for mucosal recruitment: local inflammation enables entry, while the response to cognate antigen supports the accumulation of specific MBC. Our two-step hypothesis has been recently confirmed. A recent paper found that B cell recruitment to the URT is a reactive phenomenon^74^. Seronegative volunteers were intranasally infected with SARS-CoV-2. B cells were absent at day 1 in all cases and never increased in individuals with abortive or transient infection. B cells significantly increased only in individuals with sustained infection by day 10, demonstrating that B cells reach the site of infection relatively late.

Another study investigated the induction of B-cell memory in individuals vaccinated against influenza either by the mucosal (live attenuated influenza vaccine, LAIV vaccine) or the intramuscular route (inactivated influenza vaccine, IIV)^96^. After nasal administration of the live attenuated influenza vaccine, influenza-specific B cells increased significantly in the swab by day 7, peaked at 2 weeks, and then declined. In contrast, the systemically administered inactivated vaccine had no influence on the number of MBCs in the swab, but influenza-specific MBCs increased in the peripheral blood.

These results confirm that B-cell recruitment to the swab is induced by local triggering, delayed, and short-lived^96^

Here we show that about 6% of the MBCs found in the swabs belong to large clones circulating in the peripheral blood and hypothesize that the great majority of swab MBCs may be produced by the local immune response. All MBCs, however, independently of their origin, are not found in the swab without inflammation. The mechanism of inflammation-induced B-cell recruitment may contribute to explaining why vaccine-induced memory and hybrid immunity are insufficient to prevent infection of the upper airways. Swab protection is induced after the infection is initiated and is short-lived, waning once inflammation resolves. Inflammation is the main driver of lymphocyte recruitment also to the surface of the lower airways, as suggested by the analysis of the BAL of patients with different clinical conditions.

Our results explain the phenomenon of “waning immunity” reported after vaccination against SARS-CoV-2 and other respiratory pathogens^97–99^.

Early after vaccination, a short window of strong, ready-to-use immunity is generated. Short-lived PBs secrete high levels of antibodies into the serum, which access mucosal surfaces through transudation^100,101^. ActMBCs also peak in the blood and migrate to mucosal sites when inflammation directs their homing.

Thanks to this mechanism recent vaccinations prevent infections and symptoms and can reduce virus transmission^102^. Because only about 1% of serum antibodies reach the URT^20^, this protection is fragile: once serum antibodies decline — roughly three months after the last vaccination or infection — mucosal antibody protection becomes insufficient^97,99^. Activated MBCs decay on a similar timescale, further reducing protection from prior antigen exposure. Immune memory nonetheless still protects against severe disease. Resting MBCs can be rapidly reactivated, producing high-affinity antibodies without the delay of building new germinal centers from naïve B cells. Systemic–mucosal connectivity and the mechanism of inflammation-driven homing have recently been substantiated. Following intranasal booster vaccination, systemic memory B cells were shown to migrate to the upper respiratory tract, where they underwent class switching to IgA^15,17^. B-cell recruitment was accompanied by a transient increase in cytokine and chemokine levels in nasal swabs, which are likely critical mediators of B-cell attraction^17^. Thus, the ready-to-use local protection wanes, but systemic memory is long-lived and functional.

The importance of local inflammation in the protection against respiratory pathogens has been recently confirmed by the effectiveness of mucosal vaccination with a formulation containing the TLR 4 and 7/8 ligands^103^. The vaccine, administered intranasally 3-4 times, activated innate immunity in the lungs and protected against severe infection by diverse viral and bacterial pathogens for 21 days. The addition of ovalbumin as a model antigen recruited T cells to the lung and ensured a longer lasting (3 months) broad protection against different pathogens.

In summary, our data fill a critical gap in understanding how systemic B cells contribute to immunity in URT.

Systemic mRNA vaccination, while not directly inducing BRM, primes a population of MBCs capable of migrating to mucosal sites upon local inflammation and antigen encounter. Although the contribution of systemic immunity to airway protection is relatively limited, it remains clinically significant. This is demonstrated by the effectiveness of vaccines^104–111^ and monoclonal antibodies^108,112,113^ against respiratory pathogens. At mucosal surfaces, systemic and local immune responses act in concert^11,25,63^.

We show that recruitment of MBCs to the nasal and oropharyngeal swabs is reactive rather than proactive, highlighting a fundamental limitation of systemic vaccines: they lack the capacity to pre-position MBCs at mucosal sites. On the other hand, our data suggests that BRM responses may follow the same rules, with recruitment of MBCs following rather than preceding infection. This mechanism may explain why sterilizing immunity induced by vaccination, infection, or hybrid immunity is short-lived. Protection from severe disease is always ensured by the prompt reaction of systemic and tissue-resident MBCs.

### Study Limitations

Our study has some limitations.

First, we could not compare the B cells in peripheral blood and mucosal swabs with those directly residing in the nasal lamina propria or tonsillar tissues, so the precise tissue origin of swab-detected B cells remain uncertain; they may represent a mixture of circulating, transmigrated, and tissue-resident cells.

Second, the relative low number of B cells recovered from the nasal and oropharyngeal swabs, along with the small size of the “CD27_Act_b” cluster identified in peripheral blood, limited the extent of clonal reconstruction achievable in the study.

Third, our analysis focused exclusively on CD27^pos^ B cells. We did not explore the clonal relationships between “CD27_Act_b” cells and atypical B cells population such as double-negative 2 (DN2) population (IgD^-^CD27^-^)^46^. These populations are known to increase in conditions such as autoimmunity, chronic infections, and immune deficiencies^114^. “CD27_Act_b” may correspond to the T-bet^+^ subsets that still expresses CD27^46^. Further research is needed to address this gap.

Fourth, while we demonstrate that B cells producing antibodies can be found in the nasal and oropharyngeal swabs, we could not quantify the respective contributions of these cells versus serum-derived antibodies to overall mucosal protection. This remains a critical gap with direct implications for vaccine design^10^.

Further studies are necessary to investigate whether infection modulates the relative contribution of systemic and resident B cell memory, and the transport of serum IgG to local immunity.

### Further perspective

Parenteral vaccination is known to generate long-lived B-cell immunity, and mucosal boosters may provide only transient enhancement of local response^4,115^. Our findings indicate that innovative strategies are required to more effectively prevent infection and disrupt pathogen transmission. Accordingly, we underscore the urgent need to delineate the durability of mucosal immune responses, establish robust correlates of mucosal protection, and develop platforms capable of rapidly reactivating both systemic and tissue-resident immune compartments during outbreak scenarios.

### Authors contributions

EPM and RC conceived the project. EPM, RC, and ML analyzed the data and wrote the manuscript. EPM, BLC, SB, CS, VM, EA, CA and AP perform the experiments. GV, MS and EZ performed FACS and sorting of samples. SZ, RRDP, CM and MA were responsible for patient selection. CQ, IQ, AN, CL, FL and SS revised the manuscript.

## Materials and Methods

### Resource abailability

Further information and request for resources and reagents should be directed to and will be fulfilled by the Lead Contact: Eva Piano Mortari and Rita Carsetti.

### Material and Data code Availability

All reagents will be made available on request after the completion of a Materials Transfer Agreement.

This study did not generate new unique reagents, and no novel code was created for the study.

All data are included in the manuscript and/or supporting information. The scRNA-seq and scVDJ data have been deposited in the GEO under accession number GSE (GSE281428).

### Experimental Model and Subject Details

#### Ethical statement

The study protocol was approved by the Ethics Review Committee of the Bambino Gesù Children Hospital, Rome, Italy (3194_OPBG_2023). The children’s cohort was enrolled after approval by the Ethical Review Committee of Sapienza University of Rome, Italy (Prot. 0877/ 2022). The study was conducted in accordance with the Good Clinical Practice Guidelines, the International Conference on Harmonization Guidelines, and the most recent version of the Declaration of Helsinki.

#### Human samples

This study was carried out on 144 samples obtained from 81 Health Care Workers (HCWs) enrolled at the Bambino Gesù Children’s Hospital who received the Pfizer-BioNTech (BNT162b2) mRNA vaccine. From HCWs, blood samples were obtained 9 months after 2^nd^ dose (n=11), 10 days after the 3^rd^ dose (n=11 same of T270) or 10 days after the breakthrough infection (n=15; INF). We additionally included 8 HCW samples obtained 4 years after the last SARS-CoV-2 mRNA vaccine dose, 3 HCWs sampled 10 days after the 3^rd^ dose, and 5 HCWs who received intramuscular influenza vaccine Flucelvax Tetra for which we have pre-vax sample and a sample 10 days after vaccination.

Among HCWs, we also perform nasal (n=42; n=33 with CD45^+^) and oropharynx swabs (n=49; n=24 with CD45^+^). Nasal swabs were collected from the nasal cavity epithelium approximately at the midpoint of the inferior nasal turbinate. Oropharynx swabs were performed by sampling the oropharyngeal mucosa of the posterior wall and base of the tongue. One lateral side of the oropharynx is swabbed, followed by the contralateral side. Some HCWs contributed samples at multiple timepoints and/or multiple sample types (blood, nasal swab, oropharyngeal swab), accounting for the total of 144 samples from 81 individuals. Exclusion criteria were diagnosis of primary or secondary immunodeficiency, or current use of immunosuppressive drugs.

Broncoalveolar lavage (BAL) was recovered from 11 patients with Common Variable Immunedeficiency (CVID)-associated granulomatous-lymphocytic interstitial lung disease (GLILD), from 3 patients with suspected autoimmune disease and from 1 patient with viral lung infection, as well from 2 patients with suspected lung tumor (non-inflamed controls). Children (mean age 8.6ys +/- 3.6ys) were enrolled from November 2022 to January 2023 at Policlinico Umberto I-Maternal Infantile Department in Rome. Six subjects were immunized with intramuscular vaccine Flucelvax Tetra, and six subjects were immunized with intranasal vaccine Fluenz Tetra. Two blood samples were obtained from each participant at baseline (T0), before vaccine administration, and after vaccination (T1, 20 days after vaccination). Exclusion criteria were prior influenza vaccination, diagnosis of primary or secondary immunodeficiency, ongoing infection, or current use of immunosuppressive drugs.

### Method Details

#### Cell isolation and cryopreservation

Peripheral blood mononuclear cells (PBMCs) were isolated by Ficoll Paque™ Plus 206 (Amersham PharmaciaBiotech) density gradient centrifugation and immediately frozen and stored in liquid nitrogen until use. The freezing medium contained 90% Foetal Bovine Serum (FBS) and 10% DMSO.

Nasal and oropharyngeal samples were obtained by simple localised sampling using FLOQSwab (Copan). The flocked swab was inserted into the turbinate or the upper throat and twirled 10-20 times. The swab was then placed in 1 mL of isolation medium (RPMI + 1.5mM DTT) and incubated at 37°C for 30 min. Cells were spun down at 200 x *g* for 10 min and resuspended for the flow cytometry, for the ELISpot, or for the sorting^116^.

BAL samples were collected by the surgeon and transferred to the laboratory for processing within a few hours of collection. BAL fluid was filtered through a 70 µm cell strainer and centrifuged at 1600 rpm for 10 minutes at 4°C. Cells were washed twice with RPMI medium supplemented with 2% FCS prior to staining.

### Flow cytometry and cell sorting

Flow cytometry was performed using cryopreserved PBMCs or fresh cell isolates from swabs and from BAL. Cell sorting was performed using fresh cells. Cells were stained with the appropriate combination of fluorochrome-conjugated antibodies to identify subsets of B cells according to standard techniques. Cells were acquired on a BD LSRFortessa X-20 or Symphony A5 (BD Biosciences) and sorted on an Aria II B cell (BD Biosciences). The B cell subsets were identified based on the expression of CD45, CD19, CD21, CD24, CD27, CD38, CD71, CD69, IgG, IgA, IgM, αX (CD11c), FCRL5, CXCR3, T-bet and TCF-1 by flow-cytometry. For intracellular T-bet and TCF-1 staining, cells were fixed and permeabilized with the Foxp3/Transcription Factor Staining Buffer Set (eBioscience).

Naïve B cells were identified as CD19^+^CD24^+^CD27^-^ and memory B cells were defined as CD19^+^CD24^+^CD27^+^, IgM, and switched MBCs were discriminated based on the expression of IgM, IgG or IgA. CD27^dull^ and CD27^bright^ memory B cells were discriminated and sorted due to the intensity of expression of CD27 as previously published^39^.

Plasmablasts were gated and sorted as CD19^+^CD24^-^CD38^++^. Resting MBCs were sorted as CD19^+^CD24^+^CD27^+^. Activated B cells were identified and isolated as CD71^+^ or as CD19^+^CD24^-/+^CD27^+^ after the exclusion of plasmablasts. Among actMBCs we differentiate CD27_Act_b from CD27_Act_a because of T-bet, TCF-1, CD11c and FCRL5 expression. B cells from the swabs were sorted by first excluding exfoliated epithelial cells and debris, and identifying lymphocytes based on forward- and side-scatter properties (FSC-A vs SSC-A). Subsequently, CD45□CD19□ cells were gated to define the B-cell population.

### Detection of antigen-specific B cells

For the identification of antigen-specific B cells, the biotinylated proteins were individually multimerized with fluorescently labelled streptavidin as previously described^22,23^. Separate aliquots of recombinant biotinylated antigen were mixed with streptavidin-BUV395, streptavidin-PE, streptavidin-FITC, or streptavidin-RB744 (BD Biosciences) at a 25:1 (BUV395) or 20:1 (PE, RB744, FITC) protein-to-fluorochrome molar ratio, and incubated at 4°C for 1 hour. Using this combinatorial strategy, probes were labeled as follows: SARS-CoV-2 (PE, BUV395), H1N1 HA (PE, FITC), RSV-F (BUV395, RB744), and Tetanus Toxoid (TT) (BUV395, FITC). Individually labeled antigen probes were then equimolarly combined in Brilliant Buffer (BD Biosciences) prior to use. Gating strategy was performed as previously described^117^ (Suppl. Fig. 3B), with each probe combination first gated on cells double negative for the other probe channels to exclude potential streptavidin cross-reactivity; antigen-specific B cells were then identified as double positive for binding of the same antigen labeled with its two designated fluorochromes. 4×10^6^ previously frozen PBMCs were prepared and stained with the antigen probe cocktail containing 100 ng of antigen per probe (total 200 ng). After one step of washing, surface staining with antibodies was performed in Brilliant Buffer at 4 C° for 30 min.

Stained PBMC samples were acquired on the Symphony A5 (BD Biosciences). At least 1×10^6^ cells were acquired and analyzed using Flow-Jo10.8.1 (BD Bioscience). Antigen-specific B cell phenotype analysis was performed only in subjects with at least 10 cells detected at the respective antigen-specific gate.

#### Stimulation and reagents

Sorted cells were cultured in complete medium at a concentration of 2.5×10^6^ cells/mL. Cells were then stimulated with 0.35μM CpG-B ODN2006 (Hycult Biotech); or 1 μg/mL rhCD40L (Enzo Life Sciences) plus 10 μg/mL anti-IgM/IgG/IgA (cat #109-006-064, Jackson Immunoresearch), 20 ng/mL rhIL-4 (Peprotech), and 20 ng/mL rhIL-21 (Peprotech). Complete medium was prepared as follows: RPMI-1640 (Gibco BRL, Life Technologies), 10% heat inactivated fetal bovine serum (FBS, Hyclone Laboratories Logan UT), 1% L-Glutammine (Gibco BRL); 1% Antibiotics/Antimicotics (Gibco BRL), 1% sodium pyruvate (Gibco BRL). Supernatant of nasal and oropharyngeal swabs were used to stimulate PBMCs or actMBCs alone. After culture for 1, 3 or 5 days, cells were washed, counted, and seeded to quantify antibody-secreting cells by the ELISpot assay.

#### ELISpot

ELISpot was used for the quantification of total Ig-secreting cells (IgG, IgA, and IgM) present in the PBMC samples. ELISpot MultiScreen filter plates (Millipore) were pre-wetted with 35% ethanol for ≤ 1 min and washed with ddH2O (5 × 200 µL/well). Plates were coated with affiniPure F(ab’)2 Fragment Goat anti-human IgA+IgG+IgM (H+L) (Jackson ImmunoResearch; anti-IgM concentration 2.5 µg/mL, anti-IgG 15 µg/mL, anti-IgA 10 µg/mL). Cell dilutions in RPMI were incubated in ELISpot plates for 18 h. After washing plates, secreted antibodies were detected with HRP-labelled anti-human IgG (1:2000), IgA (1:1000) or IgM (1:1000) and developed with TMB substrate (ThermoFischer) before analysis on an ELISPOT counter (Mabtech).

#### Migration assay

24-well plates with Transwell inserts (6.5-mm diameter, 5 μm pores; Corning) were used with RPMI 1640 medium (Life Technologies) supplemented with 0.5% BSA (low endotoxin; Sigma-Aldrich) as assay medium. The inserts were coated with 50 μl of human fibronectin (Sigma-Aldrich) at a concentration of 10 μg/ml in water and incubated for 1 h at 37°C in a humidified atmosphere with 5% CO₂. The solution was then aspirated and the inserts were air-dried. B cells were isolated from buffy coats using the RosetteSep Human B Cell Enrichment Cocktail (STEMCELL Technologies) followed by density-gradient centrifugation, washed, counted, and resuspended in assay medium at a final concentration of 2.5 × 10□ cells/ml. Cells were starved for 2 hours at 37°C in a humidified atmosphere with 5% CO₂. Where indicated, AMG 487 was added at a final concentration of 100 nM, 1 hour after the start of starvation. The lower Transwell chamber was filled with 600 μl of assay medium supplemented with CXCL10 (100 nM; R&D Systems), and 100 μl of cell suspension was added to the upper chamber. Cells were allowed to migrate for 3 hours at 37°C in a humidified atmosphere with 5% CO₂. Cells that migrated to the lower compartment were then collected and acquired on a Symphony A5 flow cytometer (BD Biosciences).

#### Generation of scRNA-seq and VDJ library

From peripheral blood from four healthy donors we sorted CD27^dull^, CD27^bright^, actMBC, and PBs. We sorted the different B cells among the CD27^pos^ B cells, to gather information on their ontogeny based on protein markers and not only based on mRNA. This strategy allowed us to have a better resolution in identification of rare B cell populations.

From the same four donors we obtained nasal and oropharynx swabs and sorted CD19^pos^ cells. Sorted cells were sequentially labeled using Single Cell Labeling with the BD Single-Cell Multiplexing Kit (BD Biosciences). Briefly, each donor-derived sorted population was labelled with unique sample tags, and each sample was washed twice with FACS buffer. The samples were counted on a hemocytometer (InCyto) staining the cells with Calcein AM (ThermoFisher Scientific) and Draq7^TM^ (BD Biosciences). The samples were pooled and resuspended in cold BD Sample Buffer (BD Biosciences). Single cells from the pooled sample were isolated using Single Cell Capture and cDNA Synthesis with the BD Rhapsody Express Single-Cell Analysis System following the manufacturer’s protocol (BD Biosciences). After the nanowell cartridges were primed, the pooled sample was loaded onto the BD Rhapsody cartridge and incubated at room temperature. Cell capture beads (BD Biosciences) were prepared and then loaded onto the cartridge. According to the manufacturer’s protocol, the cartridge was washed, cells were lysed, and cell capture beads were retrieved and washed before performing reverse transcription and treatment with Exonuclease I. cDNA libraries were prepared using the cDNA kit (BD Biosciences), WTA kit (BD Biosciences) and BCR kit (BD Biosciences) following the protocol 23-24018(01). The protocol allows the study of the total mRNA expression and the VDJ rearrangement for samples that have been labelled with the BD Single-Cell Multiplexing Kit. The PCR products were purified, and the mRNA PCR products, BCR products, and sample tag products were separated with double-sided size selection using Agencourt AMPure XP magnetic beads (Beckman Coulter). Total mRNA, BCR, and sample tag products were diluted to 1 ng/mL to prepare final libraries. The final libraries were indexed using PCR (9 or 6 cycles). Index PCR products were purified using magnetic beads. The quality of the final libraries was evaluated using the Agilent 2200 TapeStation with High Sensitivity D1000 and D5000 ScreenTape and quantified using a Qubit Fluorometer using the Qubit dsDNA HS Kit (ThermoFisher, #Q32854). The final libraries were diluted to 4 nM and multiplexed for paired-end (85×215) sequencing on a NovaSeq 6000 (Illumina). Both alignment, generation of barcode-gene matrices and VDJ analysis were performed through the pipeline “BD Rhapsody™ Sequence Analysis Pipeline v2.2.1” from SevenBridges (https://www.sevenbridges.com/) using as reference the human genome provided by BD (“Reference_Human_WTA_2023-02”) following the manufacturer’s recommendations. scRNA-seq data from ambient RNA was trimmed based on number of unique molecular identifiers (nUMI)-barcode saturation curve.

#### Clustering of scRNA cells and differentially expressed gene profile

The R package “*Seurat*” (v5.1.0) was used in RStudio (v4.1.5) for data trimming, unsupervised clustering, and visualisation according to the author’s guidelines^118^. Briefly, to remove poor-quality cells, we filtered cells with a gene number less than 200 and a percentage of mitochondrial genes greater than or equal to 30%. We applied a higher mitochondrial-content cutoff (30%) to retain sufficient B cells derived from the swabs for downstream analyses; importantly, retained cells displayed intact transcriptomic profiles comparable to peripheral blood B cells, indicating minimal bias from stress-related artifacts.

Each B cell library was merged with the “*harmony*” package^119^, with donor identity and tissue source specified as batch variables. Post-integration UMAPs confirmed that clusters were not driven by donor-specific effects. Previously normalisation and data scaling of the merged dataset with the “SCTransform” (flavor = v2) regressing out the mitochondrial percentage variable^118^. The first 3000 highly dispersed genes were considered highly variable genes and were used as input for principal component analysis (PCA). The first 50 PCAs were used in the subsequent analysis with a resolution of 0.5. The cells were then embedded by a uniform manifold approximation and projection for dimension reduction (UMAP) plot. To assign cell identities, we applied the “*FindAllMarkers*” to identify differentially expressed genes (DEG) among all genes by using the Wilcoxon rank sum test with standard parameters. We identified cell contaminants (less than 5% of the total cell number) that had typical macrophage, NK, and T cell markers. Therefore, these cells were removed from the analysis and the integration protocol was repeated, using the same settings with a different resolution parameter of 0.2.

For the definition of the B cell clusters, we used expert-curated knowledge and gene signatures derived from previously published literature^33,39^. Briefly, to infer the CD27 cluster identities, we applied the following signatures (CD27^dull^: TCL1a, BTLA, BACH2, CCR7, CXCR4, IL4R. CD27^bright^: TLR9, CXCR3, ITGAM, CD27, IKZF1, TBX21, GAPDH, POU2AF1. ActMBCs: MS4A1, CD52, CD86, CD1C, CXCR3, HLA-DRA, TFRC. Plasmablast: XBP1, CD38, PRDM1, JCHAIN, IRF4) with the “AddModuleScore” function with a number of control features (*ctrl*) set to 15. To integrate the data from the gene signatures, we considered the overall percentage of each cluster distribution across the sorted populations (CD27^dull^, CD27^bright^, actMBCs, PBs, and mucosal B cells).

#### Single-cell VDJ analysis

Single-cell VDJ clonotype analysis was performed using the AIRR output of the “BD Rhapsody™ Sequence Analysis Pipeline”. We used the R package “*imncantation*” (v4.5.0) framework to build phylogenetic tree on BCR full length data following the author’s recommendations. Briefly, after the removal of cells having multiple or no heavy chains, we analyzed only cells that were matched between the scVDJ and scRNA-seq data sets. With the ‘SCOPer’ R package (v1.3.0) on IGH we determined the clonal clustering threshold (the threshold was found using the ‘findThreshold’ function with standard parameters), assigned clonal groups and reconstruct germline sequences. For the germline, we used a reference database of known alleles of IMGT (https://www.imgt.org/). Mutational frequency was performed using the “SHazaM” R package (v1.2.0) “observedMutations” function using as subregion for mutation calculation the parameter “regionDefinition” = IMGT_VDJ_BY_REGIONS, where CDR1, CDR2, CDR3, FWR1, FWR2, FWR3 and FWR4 regions are treated as individual regions. The mutation frequencies were based on the nucleotide sequences.

We finally built and visualized trees with the “Dowser” R package (v2.2.0) using the maximum likelihood trees from phangorn. The trees were visualised using the R package ‘ggTree’ (v3.12.0).

### scRNA-seq analysis of published data from nasopharyngeal swabs of SARS-CoV-2 infected individuals

The processed scRNA-seq object of the nasopharyngeal swabs from Lindeboom et al., 2024 were retrieved from the COVID-19 Cell Atlas web portal (https://covid19cellatlas.org). Briefly, the H5AD file was converted into a Seurat object and explored in Seurat previously normalization, scaling and integration with Harmony using the sequencing library (“sequencing_library”) as variable and the first 30 PCA dimensions. The author cluster annotation was adopted. B cells were filtered and cell contaminants were removed by the lack of known B cell markers *MS4A1*, *CD19*, *CD79A*.

### Statistical methods

Data were first tested for normality and homoscedasticity using Shapiro Wilk and Levene tests, and since assumptions were violated, nonparametric tests were used for analysis. The Wilcoxon matched pair signed-rank test or the two-tailed Mann–Whitney U-test were used. Categorical variables were compared using the exact Chi-square test. A two-sided p value less than 0.05 was considered statistically significant. Linear regression analysis was used to assess the relationship between the mean fluorescence intensity (MFI) and the frequency (%). Pearson correlation analysis was performed to quantify the strength of the association. All statistical analyses were performed with GraphPad Prism 10.2.3 (GraphPad Software).

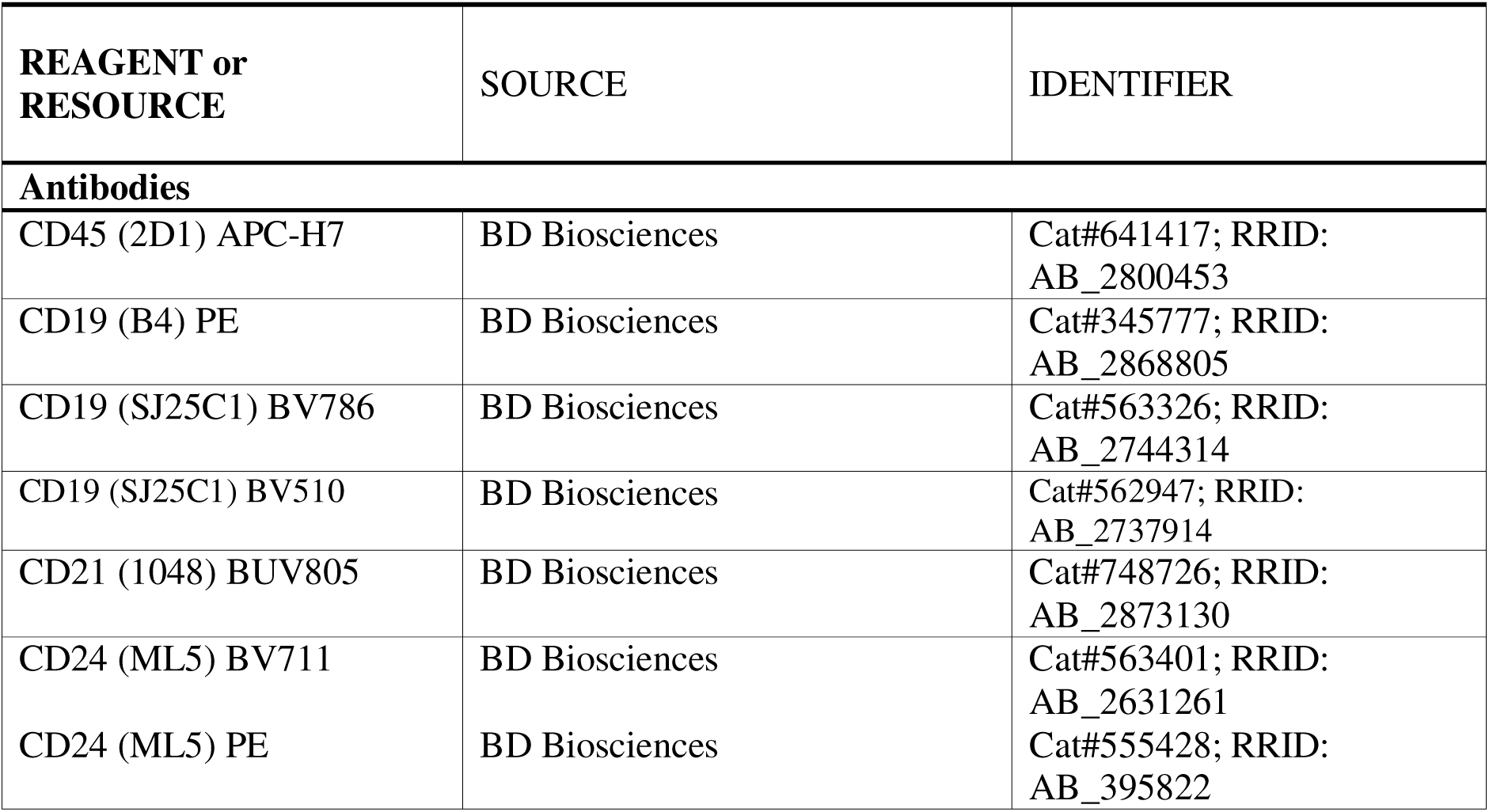
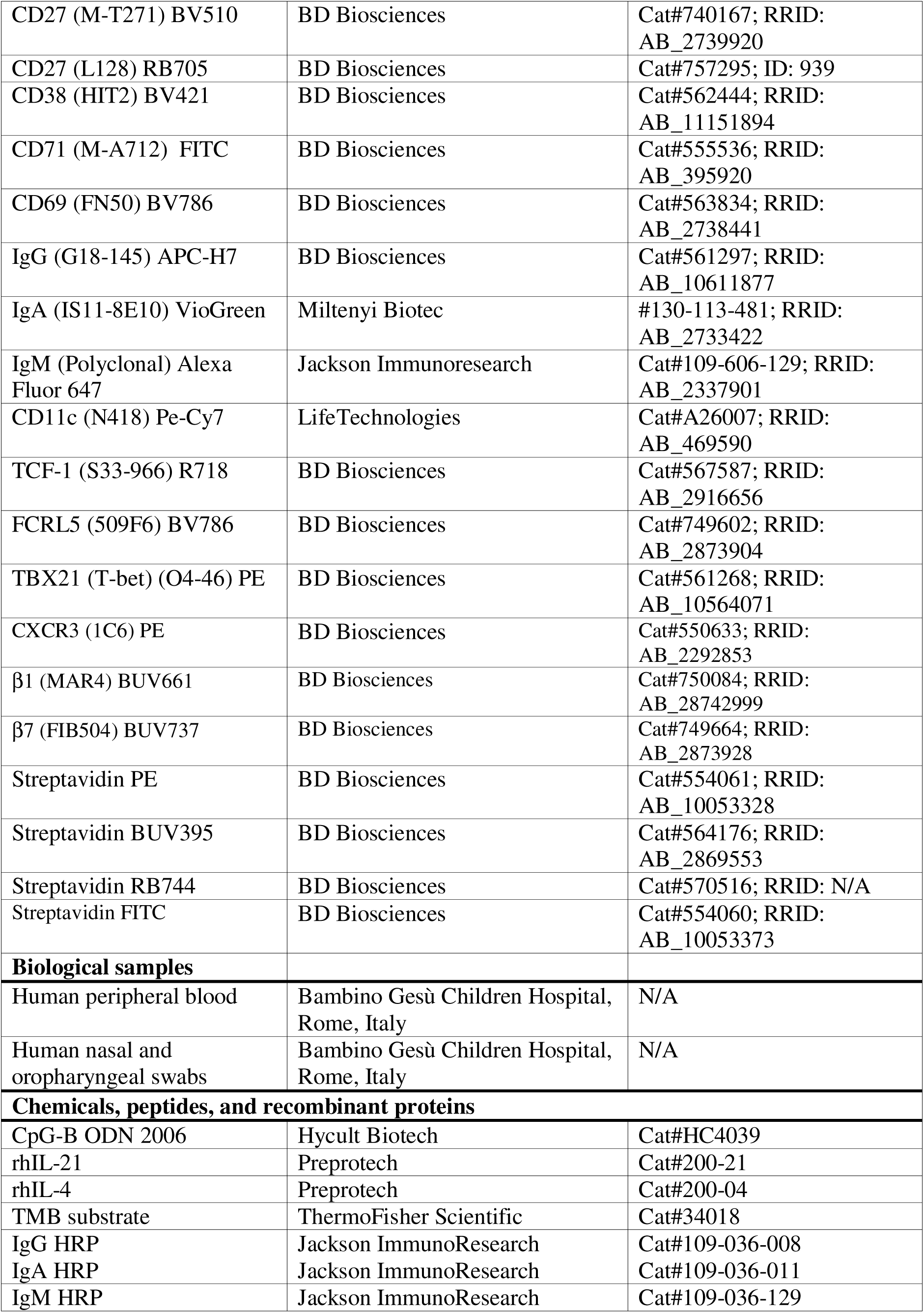
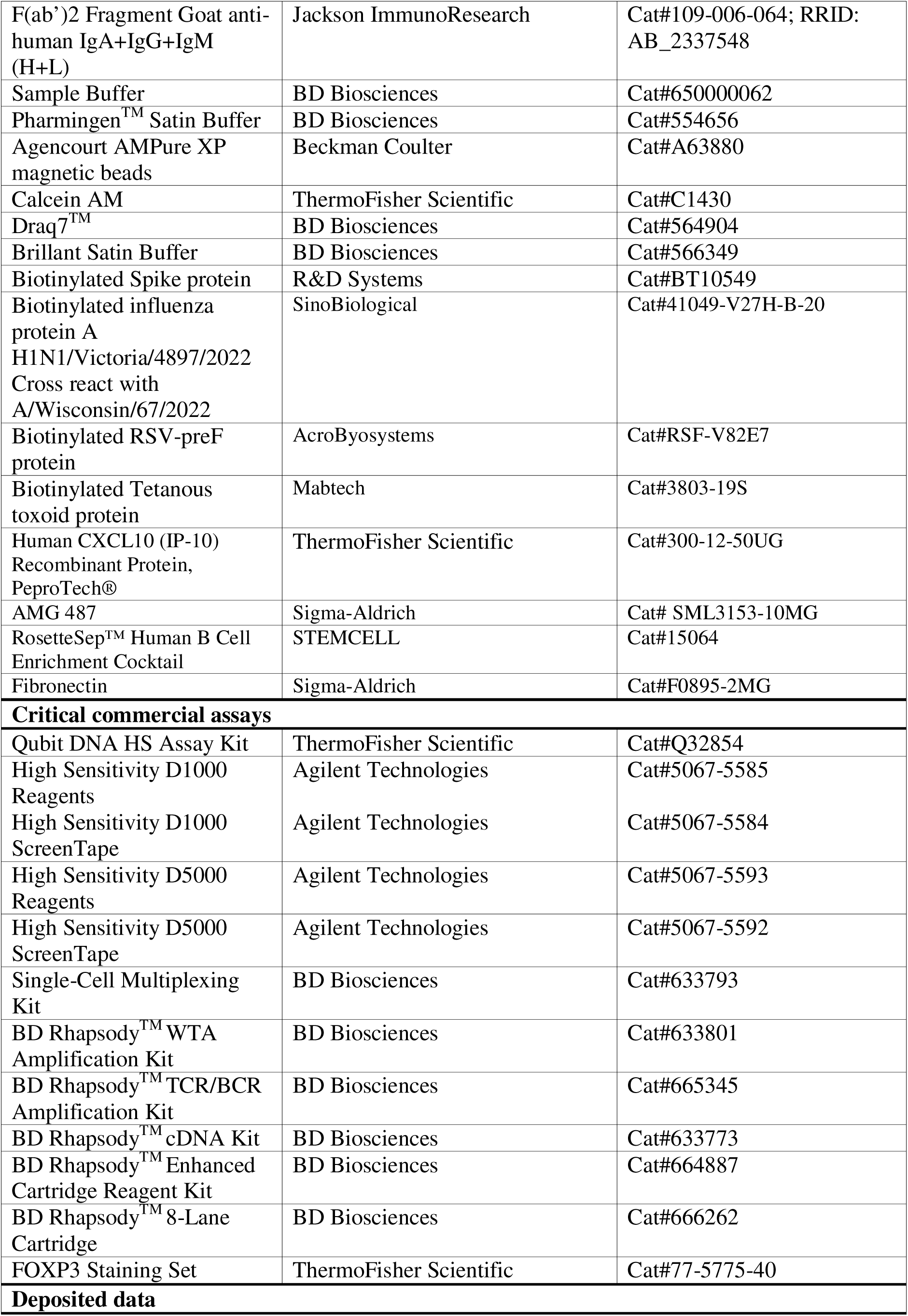
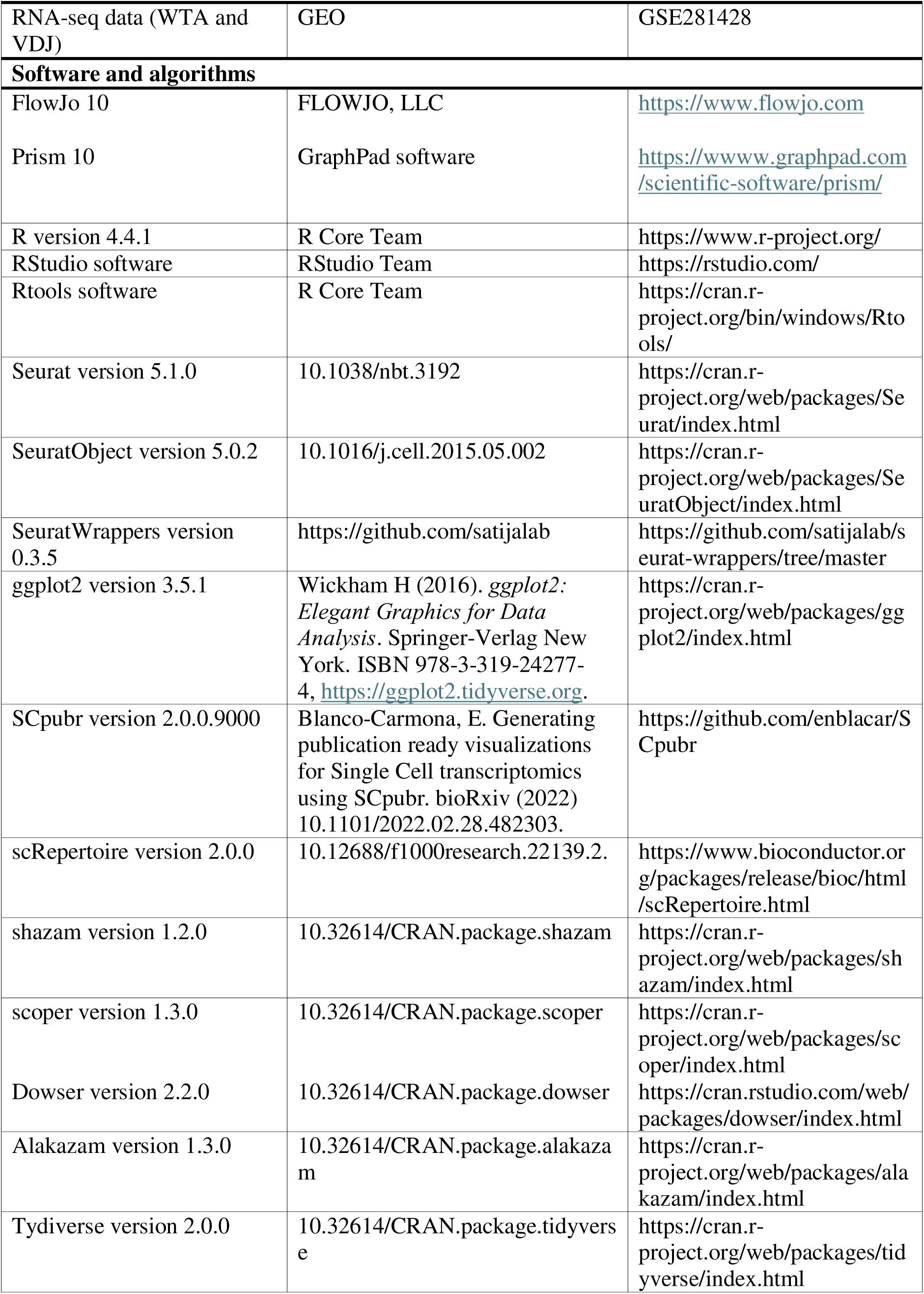
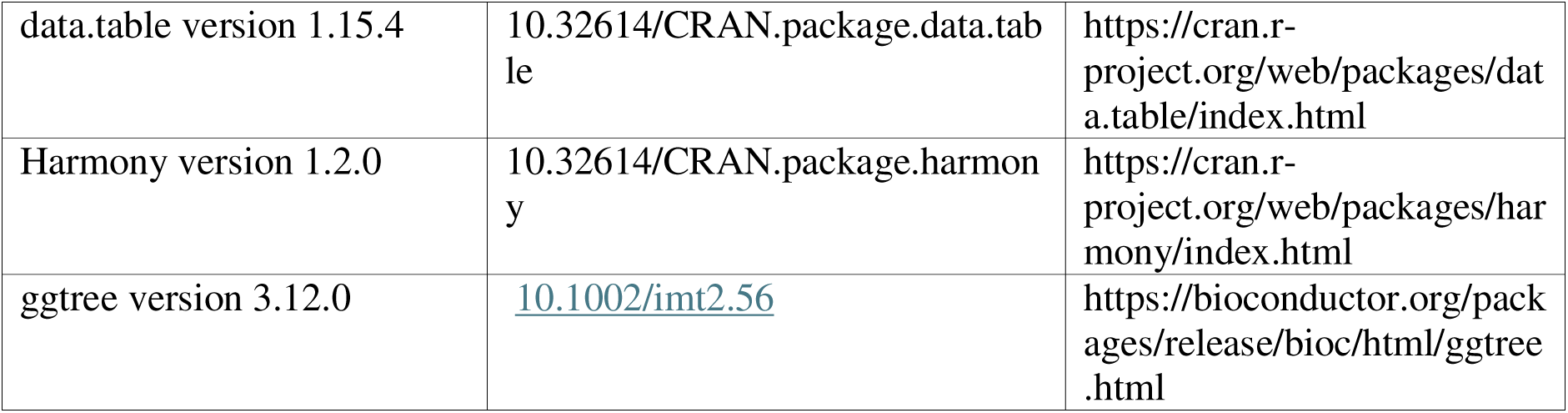

## Acknowledgments

EPM was supported by ESCMID-European Society of Clinical Microbiology and Infectious Diseases (202303_ESCMID_MORTARI) and also by the Italian Ministry of Health with Current Research funds (Ricerca Corrente and 5xmille).

SS was supported by the Italian Ministry of Health with the grant COVID-2020-12371735; and the PRIN-2022 (20228KZKE3).

The research leading to these results has received funding from the European Union - NextGenerationEU through the Italian Ministry of University and Research under PNRR - M4C2-I1.3 Project PE_00000019 “HEAL ITALIA” to IQ CUP: B53c22004000006.

We thank Manuela Colucci from the Bambino Gesù Children’s Hospital and Irene Mattiola from the Charitè –Universitätsmedizin Berlin for kindly reviewing this paper and offering helpful comments.

## Supplementary Materials

**Suppl. Figure 1:**
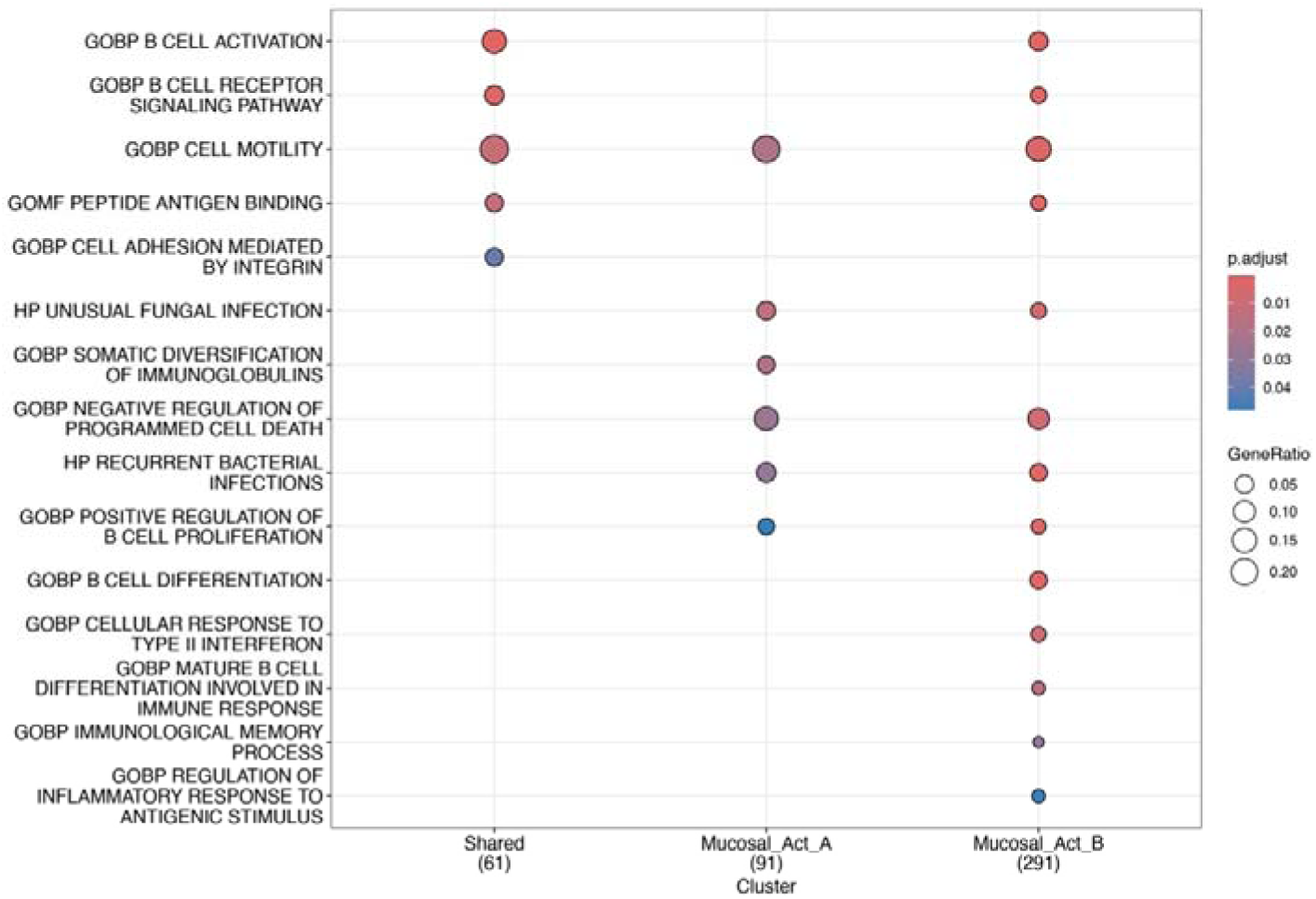
Gene set enrichment analysis (GSEA) of biological pathways shared across or specifically enriched within B_Mucosal, CD27_Act_a, and CD27_Act_b clusters.

**Suppl. Figure 2:**
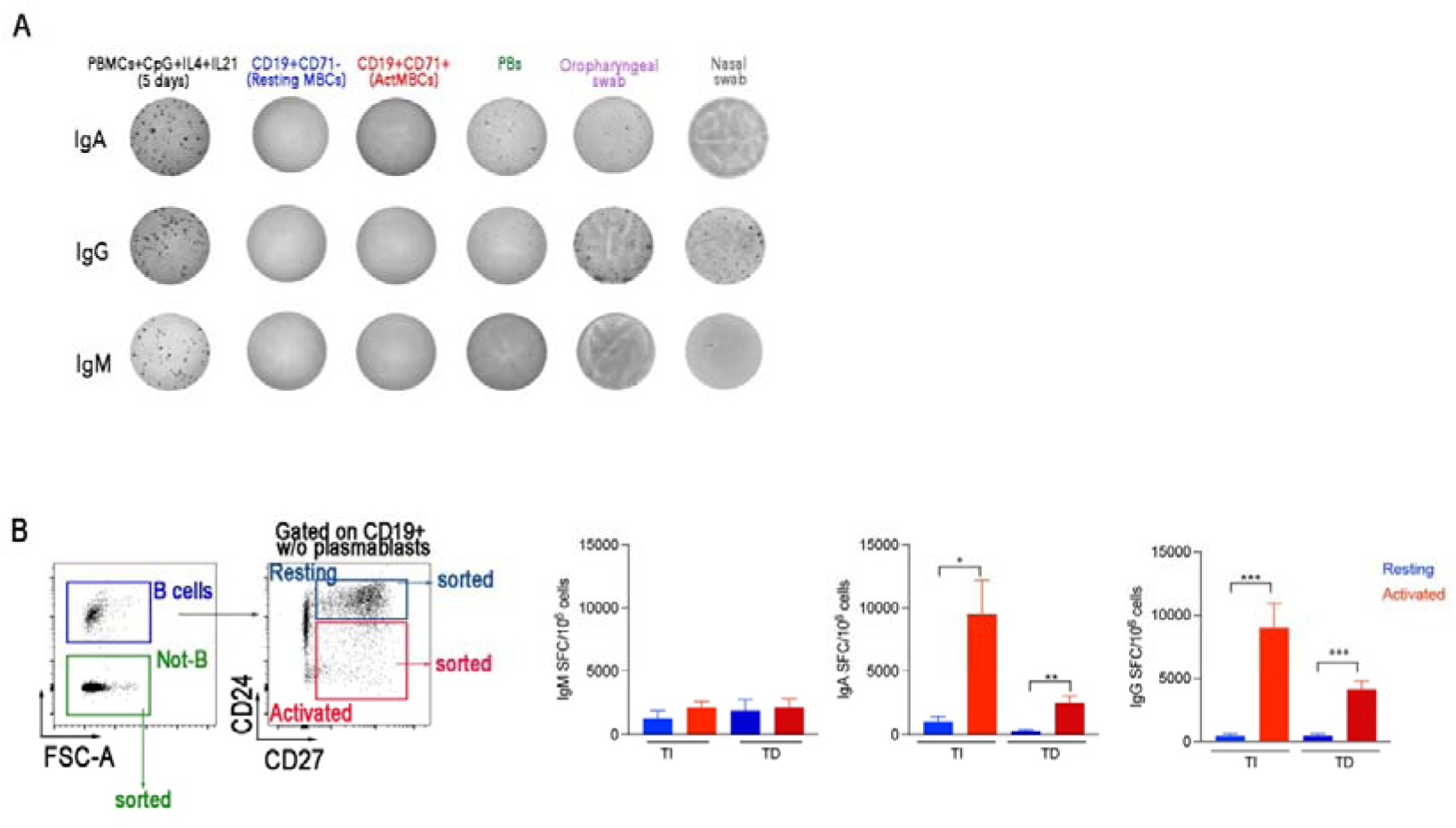
(A) Representative ELISpot (total IgA, IgG, and IgM) obtained by plating total PBMCs stimulated for 5 days with CpG, IL-21, and IL-4, non-stimulated actMBCs (sorted as CD71^+^), resting B cells (sorted as CD71^-^) and PBs (sorted as CD24^-^CD38^++^). B cells isolated from the nasal and oropharyngeal swabs were plated directly without previous stimulation. (B) Gating strategy to sort, not-B cells, resting and activated MBCs. Total IgA, IgG, and IgM obtained by plating not-B cells + actMBCs or not-B cells + resting MBCs stimulated for 3 days with TI or TD stimuli. Columns depict mean + SEM. The nonparametric Mann-Whitney t test was used to evaluate statistical significance. The significances of the two-tailed P value are shown as *p< 0.05, ***p<0.001.

**Suppl. Figure 3:**
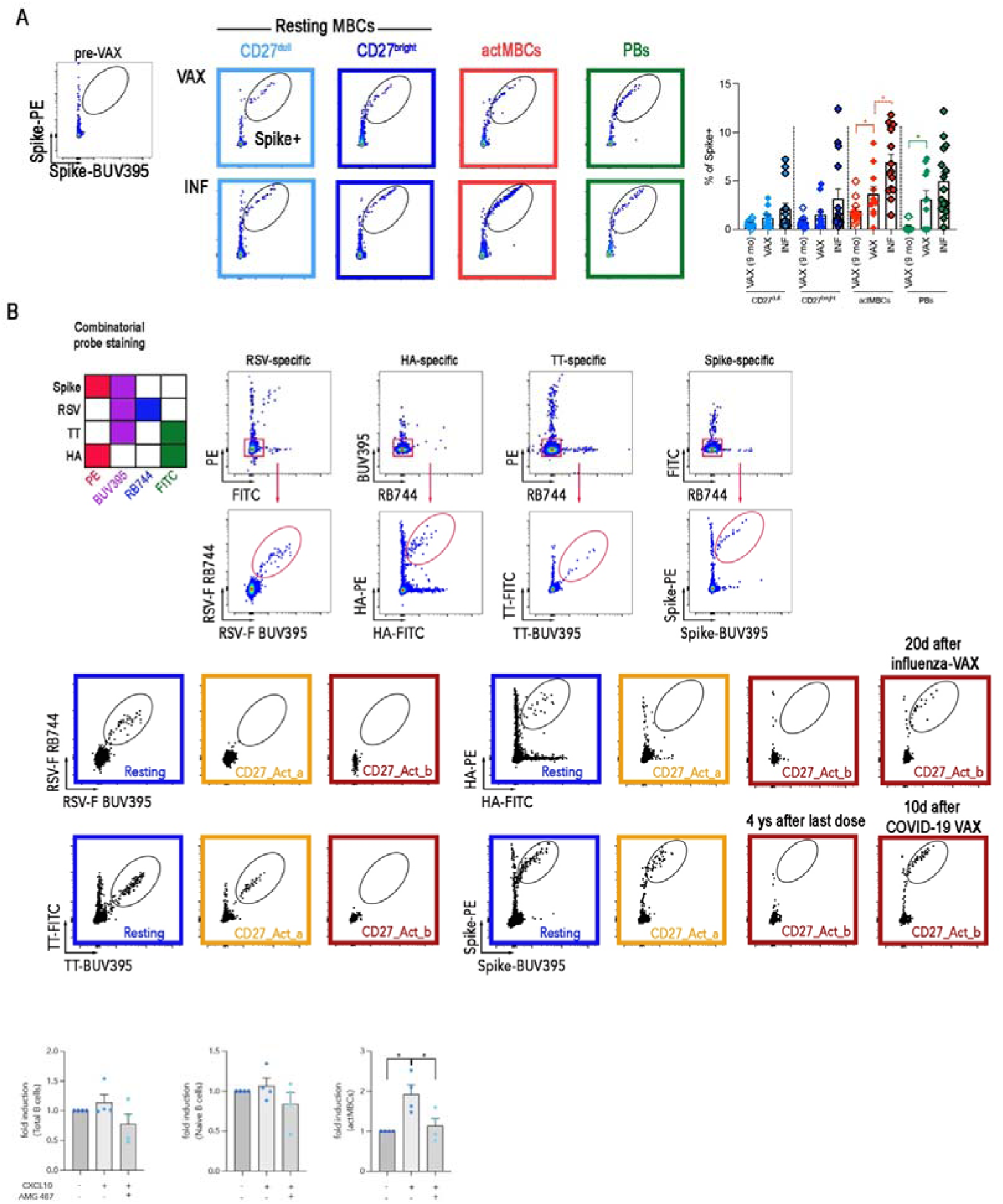
(A) Identification of Spike-specific B cells among CD27^dull^, CD27^bright^, actMBCs and PBs in two representative adult subjects (post 3^rd^ dose: VAX, and 10 days after breakthrough infection: INF). The graph illustrates the frequency of Spike-specific resting MBCs (CD27^dull^ and CD27^bright^), actMBCs and PBs among all subjects analyzed 9 months after the 2^nd^ dose, 10 days after the 3^rd^ dose or after a breakthrough infection. A representative pre-pandemic FACS plot is also shown. (B) Combinatorial probe staining and gating strategy for detecting multiple B cell specificities. To exclude B cells potentially cross-reactive to streptavidin, each probe combination was first gated on cells negative for the other two probe channels. Antigen-specific B cells were then identified as double positive for binding to the same antigen multimerized with two distinct fluorochromes, following a matrix coding scheme. Representative FACS plot for the identification of RSV-, Flu-, TT- and Spike-specific B cells among resting, CD27_Act_a and CD27_Act_b MBCs. (C) Bar graphs display the fold induction of migration relative to spontaneous migration (set to 1) for total B cells, naïve B cells and total actMBCs. Columns depict mean + SEM. The nonparametric Wilcoxon pair signed rank test (continuous line) and Mann-Whitney t test (dotted line) were used to evaluate statistical significance. The significances of the two-tailed P value are shown as *p< 0.05, **p<0.01, ***p<0.001, ****p<0.0001.

**Suppl. Table 1.excel**

**Suppl. Table 2.excel**

**Suppl. Data.pdf**

